# Arabidopsis mTERF9 protein promotes chloroplast ribosomal assembly and translation by establishing ribonucleoprotein interactions *in vivo*

**DOI:** 10.1101/2020.06.16.153288

**Authors:** Louis-Valentin Méteignier, Rabea Ghandour, Aude Zimmerman, Lauriane Kuhn, Jörg Meurer, Reimo Zoschke, Kamel Hammani

## Abstract

The mitochondrial transcription termination factor proteins are nuclear-encoded nucleic acid binders defined by degenerate tandem helical-repeats of ~30 amino acids. They are found in metazoans and plants where they localize to mitochondria or chloroplasts. In higher plants, the mTERF family comprises ~30 members and several of these have been linked to plant development and response to abiotic stress. However, knowledge of the molecular basis underlying these physiological effects is scarce. We show that the Arabidopsis mTERF9 protein promotes the accumulation of the *16S* and *23S* rRNAs in chloroplasts, and interacts predominantly with the *16S* rRNA *in vivo* and *in vitro*. Furthermore, mTERF9 is found in large complexes containing ribosomes and polysomes in chloroplasts. The comprehensive analysis of mTERF9 *in vivo* protein interactome identified many subunits of the 70S ribosome whose assembly is compromised in the null *mterf9* mutant, putative ribosome biogenesis factors and CPN60 chaperonins. Protein interaction assays in yeast revealed that mTERF9 directly interact with these proteins. Our data demonstrate that mTERF9 integrates protein-protein and protein-RNA interactions to promote chloroplast ribosomal assembly and translation. Besides extending our knowledge of mTERF functional repertoire in plants, these findings provide an important insight into the chloroplast ribosome biogenesis.

## INTRODUCTION

The mitochondrial transcription termination factor (mTERF) proteins are tandem degenerate α-helical repeats proteins that are encoded by nuclear genomes of all eukaryotes except fungi (1). The mTERF family was named for its founding member, a human mitochondrial protein that promotes transcription termination *in vitro* (2). Each mTERF repeat spans ~30 amino acids that fold into two consecutive antiparallel α-helices followed by a shorter α-helix perpendicular to the first one (3–5). The mTERF repeats stack together to form an elongated solenoid structure with a central groove capable of binding nucleic acids (5). mTERF proteins typically harbor an N-terminal organellar transit peptide and localize to mitochondria or chloroplasts and are considered to be putative organellar gene regulators (reviewed in 6). Whereas metazoans have 3 to 4 mTERF members, some plant genomes encode more than 30 mTERF proteins (1,7,8). The functions of mTERF proteins were first characterized in metazoans showing that they influence mitochondrial gene transcription, DNA replication and ribosome biogenesis (reviewed in 9,10). In plants, several of these genes are essential for embryo viability (11–13). Others have been linked to a variety of abiotic stress-responses (8,14–17) but how these genes trigger these responses in plants is not understood. Plant mTERFs are predicted to act in mitochondria or chloroplasts but knowledge about their roles in organelles is scarce. In fact, only five of the ~30 mTERF proteins found in angiosperms have been connected to their gene targets and functions in organelles. In Arabidopsis, mTERF5 (also known as MDA1), mTERF6 and mTERF8 are chloroplast DNA binding proteins involved in the regulation of chloroplast gene transcription (18–20). mTERF5 stimulates the initiation of transcription of the *psbE* and *ndhA* genes (18,21), whereas mTERF8 and mTERF6 promote the termination of transcription of *psbJ* and *rpoA*, respectively (19,20). mTERF6 has additionally been reported to affect the maturation of *trnI.2* but the reason for this effect remained unclear (22). Finally, mTERF15 and mTERF4 contribute to the RNA splicing of the *nad2-3* intron in Arabidopsis mitochondria and group II introns in maize chloroplasts, respectively. Therefore, to date the functional repertoire of mTERFs in plant organelles concerns the regulation of gene transcription and intron splicing. In Arabidopsis, the *mTERF9* gene (known as well as *TWIRT1*) encodes a chloroplastic mTERF protein that has been involved in the development of the shoot apical meristem (23) and the plant acclimation to high salinity (17,24) and photo-oxidative stress (25). However, the function of mTERF9 in chloroplasts was not further studied and the molecular basis underlying its physiological effects on plants is unknown. To answer this question, we examined the molecular defects in the *mterf9* mutant and characterized the primary functions of mTERF9 in Arabidopsis. We show that mTERF9 is required for chloroplast ribosomal assembly and therefore, translation. We performed a comprehensive analysis of the RNA and proteins bound by mTERF9 *in vivo* and demonstrated its predominant interaction with the *16S* rRNA and a large set of proteins required for the biogenesis of the small ribosomal subunit in chloroplasts. Our findings further reveal that mTERF9 can support direct interactions with both protein and RNA ligands which likely account for the protein function in the ribosomal assembly *in vivo*. Finally, we demonstrated that mTERF9 interacts physically with the CPN60 chaperonin complex *in vivo* suggesting a functional cooperation between these proteins in the chloroplast ribosome biogenesis and translation. This work expands the functional repertoire ascribed to plant mTERF proteins in translation and provides mechanistic insights into their *in vivo* functions in organellar gene expression.

## MATERIALS AND METHODS

Oligonucleotides used in this study are listed in Supplementary Table 1.

### Plant material

*Arabidopsis thaliana* ecotype Columbia (Col-0) and *Nicotania benthamiana* were used in this study. The T-DNA insertion mutant allele *mterf9* (WiscDsLox474E07) was obtained from the ABRC Stock Center. Complemented mutants were obtained via *Agrobacterium tumefaciens* transformation of *mterf9* homozygous plants. The binary vector (pGWB17) used for agro-transformation expressed the At5g55580 coding sequence in fusion with a 4xMyc C-terminal tag under the control of the CaMV 35S promoter. Transgenic plants were selected on Murashige and Skoog (MS) plates containing 25 μg/mL hygromycin. Experiments were performed using 7-day-old plants grown *in vitro* (1× MS pH5.7, 0.5% sucrose, 0.8% Agar; 16 h light: 8 h dark cycles; 65-85 μmol photons m^−2^ s^−1^), 14-day-old plants grown on soil for chloroplast isolation or 4-week-old plants for protein pulse labelling experiments.

### Subcellular localization of mTERF9

*Nicotiana benthamiana* leaves were infiltrated with *Agrobacterium tumefaciens* GV3101 carrying pMDC83*:mTERF9* and pB7RWG2*:RAP* at an OD_600_ of 0.5 each. Protoplasts were prepared as described previously (26) and examined under a Zeiss LSM 780 confocal microscope. GFP was excited at 488 nm and emission was acquired between 493-556 nm. RFP and chlorophyll were excited at 561 nm and emissions were acquired between 588-641 nm and 671-754 nm, respectively.

### Chlorophyll a fluorescence induction and light-Induced PSI absorbance changes

Chlorophyll *a* fluorescence induction kinetics and PSI absorbance changes at 820 nm were performed with leaves of 2-week-old WT, *mterf9* mutant and complemented mutant plants grown on soil using a Dual-PAM-100 System (Walz, Effeltrich, Germany) (27). Φ_PSI_, Φ_PSI NA_ and Φ_PSI ND_ were expressed as described (28).

### RNA analyses

Tissues were ground in liquid nitrogen and RNA was extracted with Trizol following manufacturer’s protocol (Invitrogen™). RNA was further extracted with phenol-chloroform pH 4.3. Five μg of Turbo DNase (Thermo Fisher) treated RNAs were used for Superscript IV reverse transcription with random hexamers. The resulting cDNA was diluted 20-fold for qPCR reaction. *ACT2 (AT3G18780)* and *TIP41 (AT1G13440)* were used as reference genes. For rRNA gel blotting, 0.5-1 μg of RNA (10 μg for other transcripts) was fractionated on 1.2% agarose-1% formaldehyde gel and blotted as described (29). Gene PCR products of 200-300 bp were labelled with ^32^P-dCTP following the prime-a-gene labelling kit instructions (Promega) and used as probes (Supplementary Table 1). Results were visualized on an Amersham Typhoon imager and data quantification was performed with ImageJ.

### Protein analyses

For *in vivo* labeling of chloroplast proteins, leaf discs of *Arabidopsis* plants were incubated in 1 mM K_2_HPO_4_/KH_2_PO_4_ pH 6.3, 1% Tween-20, 20 μg/mL cycloheximide, 100 μCi ^35^S-methionine and vacuum infiltrated. Leaf discs were kept under light for 15 min, washed in water and frozen in liquid nitrogen. Proteins were extracted in Tris pH 7.5, 10% glycerol, 1% NP40, 5 mM EDTA, 2 mM EGTA, 35 mM β-mercaptoethanol, 1× EDTA-free protease inhibitor cocktail (Roche), and 200,000 cpm per sample were resolved on SDS-PAGE. After electrophoresis, the gel was stained in 50% methanol, 10% glacial acetic acid, 0.5 g/l Coomassie brilliant blue R-250 and vacuum dried before being exposed to a phosphorimager plate. Results were visualized on an Amersham Typhoon imager. For immunoblot analysis, total leaf proteins were extracted in the same buffer, resolved on SDS-PAGE and transferred onto PVDF membrane at 80 V for 1,5 h using the wet transfer. Anti-PsaD, -PetD and -RH3 antibodies were donations of Alice Barkan (University of Oregon). Anti-NdhL, -NdhB and -RbcL antibodies were donations of Toshiharu Shikanai (University of Kyoto) and Géraldine Bonnard (CNRS UPR2357), respectively. Other antibodies against chloroplast proteins were purchased from Agrisera and anti-Myc antibodies (clone 9E10) from Sigma-Aldrich.

### Chloroplast isolation and fractionation

Chloroplasts were purified by density gradient and differential centrifugations as described previously (30). Chloroplasts were lysed in 30 mM Hepes-KOH pH 8, 10 mM Mg(OAc)_2_, 60 mM KOAc, 1 mM DTT, 1× EDTA-free protease inhibitor cocktail and 1 mM PMSF. Stromal (soluble) and thylakoid proteins were separated by centrifugation at 20,000 g for 10 min at 4°C.

### Sucrose gradient fractionation

For the analysis of high-molecular-weight complexes by differential sedimentation, 0.25 mg of stromal proteins were fractionated on 10-30% linear sucrose gradient at 235,000 g for 4 hours at 4°C as described (31). Proteins from each fraction were ethanol precipitated overnight at 4°C before their fractionation on SDS-PAGE. Polysome analyses were performed on leaf tissues as described (29). Briefly, 0.4 mg of leaf tissue was ground in 1 mL of cold polysome extraction buffer (200 mM Tris pH 9, 200 mM KCl, 35 mM MgCl_2_, 25 mM EGTA, 200 mM sucrose, 1% triton X-100, 2% polyoxyethylene-10-tridecyl ether, heparin 0.5 mg.mL^−1^, 100 mM β-mercaptoethanol, 100 μg. mL^−1^ chloramphenicol, 25 μg. mL^−1^ cycloheximide) and the extract was cleared by filtration and centrifugation. Polysomes were treated or not with 500 μg. mL^−1^ puromycin / 500 mM KCl at 37°C for 10 minutes before adding 0.5% sodium deoxycholate. Insoluble material was pelleted at 16,000 g for 10 min at 4°C and soluble extracts were fractionated on linear 15-55% sucrose gradient at 235,000g for 65 min at 4°C. For RNA isolation, 200 μL of sucrose gradient fraction was mixed with 400 μL of 8M Guanidine-HCl to dissociate RNPs and RNAs were precipitated by the addition of 600 μL ethanol 100% and incubation at −20 °C overnight. Proteins were precipitated as described above.

### CoIP-MS

Two mg of stromal proteins treated or not with 100 μg/mL RNase A and 250 U/mL RNase T1 mix (Thermo Fisher) were diluted in one volume of Co-IP buffer (20 mM Tris pH 7.5, 150 mM NaCl, 15 mM MgCl_2_, 0.5 mM DTT, 3mM ATP, 1 mM EDTA, 1% NP-40, 1× EDTA-free protease inhibitor cocktail, 1 mM PMSF) and incubated with 50 μL of anti-Myc Miltenyi magnetic beads at 4°C for 30 min on a rotator. Beads were washed in Co-IP buffer and eluted as recommended by the manufacturer. Eluted proteins were prepared as described (21,32). Briefly, proteins were precipitated overnight with 5 volumes of cold 0.1 M ammonium acetate in 100% methanol and digested with sequencing-grade trypsin (Promega) and each sample was analyzed by nanoLC-MS/MS on a QExactive+ mass spectrometer coupled to an EASY-nanoLC-1000 (Thermo Fisher). Data were searched against the *Arabidopsis thaliana* TAIR database with a decoy strategy (release TAIRv10, 27282 forward protein sequences). Peptides and proteins were identified with Mascot algorithm (version 2.5.1, Matrix Science, London, UK) and data were further imported into Proline v1.4 software (http://proline.profiproteomics.fr/). Proteins were validated on Mascot pretty rank equal to 1, and 1% FDR on both peptide spectrum matches (PSM score) and protein sets (Protein Set score). The total number of MS/MS fragmentation spectra was used to relatively quantify each protein (Spectral Count relative quantification). Proline was further used to align the Spectral Count values across all samples. The mass spectrometric data were deposited to the ProteomeXchange Consortium via the PRIDE partner repository (33) with the dataset identifier PXD018987 and 10.6019/PXD018987.

For the statistical analysis of the co-immunoprecipitation proteomes, the mass-spectrometry data collected from three biological replicates of the experimental mTERF-Myc coIPs were compared to biological triplicates of control WT coIPs using RStudio v1.1.456 and the R package IPinquiry v1.2. The size factors used to scale samples were calculated according to the DESeq2 normalization method (34). EdgeR v3.14.0 and Stats v3.3.1 were used to perform a negative binomial test and calculate the fold changes and adjusted *P*-values corrected by Benjamini–Hochberg for each identified protein. The −log_10_ (adj_*P*) and volcano plot graphs were calculated and drawn with Excel, respectively. The functional protein annotations were retrieved from the TAIR database (35) using the bulk data retrieval tool. The complete list of protein interactants and the number of peptides are provided in Supplementary Data Set 1.

### Yeast two hybrid analysis

Coding sequences of *mTERF9* or putative interacting partners were cloned into the bait vector pDHB1 or prey vector pPR3-N (Dualsystems Biotech) (36). The NMYZ51 yeast strain was co-transformed with bait and prey vectors using the PEG/LiOAc method (37). Co-transformants were selected on yeast synthetic and drop-out (DO) minus leucine (L) and tryptophan (W) agar medium. Positive colonies were sub-cultured in -WL DO liquid medium overnight. Overnight cultures were diluted to an OD_600_ of 0.3 to make the starting cultures and diluted by tenfold to 10^−2^. Five μL of each dilution was plated on -WL DO agar medium or on DO medium minus leucine, tryptophan, histidine and adenine (-WLHA) supplemented with 3-aminotriazol to select protein interactions. 3-AT was used at concentrations of 1 mM to test mTERF9 interaction with ERA1 and mTERF9, 2 mM with PSRP2 and RPL1 and 40 mM for CPNB1 and CPNB3. The expression of bait and prey proteins in yeast were confirmed by immunoblotting on total yeast protein extracts. 5 mL of saturated yeast culture (OD_600_=3) was centrifuged at 800 g for 5 min. The pellet was resuspended in 200 μL of 2 M NaOH and incubated 10 min on ice. One volume of 50% TCA was added and the mixture was incubated for 2 hours on ice and centrifuged 20 min at 16,000g at 4°C. The pellet was dissolved in 200 μl of 5% SDS before adding 200 μL of protein loading buffer (25 mM Tris pH 6.8, 8 M UREA, 1 mM EDTA, 1% SDS, 700 mM β-mercaptoethanol, 10% glycerol) and incubated at 37°C for 15 min under agitation. Extracts were cleared by centrifugation for 5 min at 16,000g at 20°C and the supernatant fractions were kept for immunoblot analysis. Ten μL of protein fractions were resolved by SDS-PAGE and transferred to a PVDF membrane. Bait and prey proteins were immunodetected using antibodies against LexA and HA antibodies respectively, purchased from Sigma-Aldrich.

### RNA immunopurification analysis

0.5 mg of stromal proteins were diluted in 450 μL of RIP buffer (20 mM Tris pH 7.5, 150 mM NaCl, 1 mM EDTA, 1% NP-40, 1× EDTA-free protease inhibitor cocktail, 1 mM PMSF) and incubated with 50 μL of anti-MYC Miltenyi magnetic beads at 4°C for 30 min on a rotator. Beads were washed and eluted in RIP buffer supplemented with 1% SDS. Immunoprecipitated and supernatant RNAs were extracted with Trizol and further purified with phenol/chloroform. The RNA from the pellet and 3.5 μg RNA from the supernatant were fragmented and labelled with Cy5 (635 nm) and Cy3 (532 nm), respectively and hybridized on a tilling microarray (chip) covering the Arabidopsis chloroplast genome, as described in (38). Data were analyzed with GenePix Pro 7.0 software with local background subtraction method. The median of ratios of the background-subtracted pellet to supernatant signals were calculated and the super-ratios of the mTERF9 IP to control IP were plotted along the Arabidopsis chloroplast genome. RIP-chip data are provided in Supplementary Data Set 2. For qRT-PCR analysis, half of the input and IP RNAs were treated with Turbo DNase (Thermo Fisher) and cDNA synthesis and qPCR were conducted as described above.

### Expression of recombinant mTERF9

The *mTERF9* sequence coding for the mature mTERF9 (amino acids 45 to 496) lacking the chloroplast transit peptide was amplified by PCR on Arabidopsis cDNA and cloned into pMAL-TEV vector within BamHI and SalI restriction sites. The N-terminal MBP fusion protein (rmTERF9) was expressed in *E. coli* and purified by amylose affinity chromatography as described (21). The purity of the recombinant protein was visualized on SDS-PAGE and Coomassie Brilliant Blue staining. The band migrating at the expected size of rmTERF9 (~96 kDa) and a comigrating band (~60 kDa) were gel excised and analyzed by mass spectrometry (LC-MS/MS) to confirm their identity.

### Northwestern blot analysis

Recombinant proteins were electrophoresed on a SDS-10% polyacrylamide gel and electroblotted to a PVDF membrane. After transfer, proteins were renatured by incubation of the membrane overnight at 4°C in renaturation buffer (100 mM Tris pH 7.5, 0.1% NP-40). Membranes were subsequently blocked for 10 minutes at 23°C in blocking buffer (10 mM Tris pH7.5, 5 mM Mg(OAc)_2_, 2 mM DTT, 5% BSA, 0.01% Triton X-100). Blocked membranes were hybridized for 4 hours at 4°C in 5 mL of hybridization buffer (10 mM Tris pH7.5, 5 mM Mg(OAc)_2_, 2 mM DTT, 0.01% Triton X-100) containing 0.5 or 1 fmole of [α-^32^P]-labelled RNA probes. Membranes were washed 4 times in wash buffer (10 mM Tris pH7.5, 5 mM Mg(OAc)_2_, 2 mM DTT) and exposed to a phosphorimager plate. Results were visualized on an Amersham Typhoon imager.

### Accession numbers

The gene described in this article corresponds to the following Arabidopsis Genome Initiative code: At5g55580 (mTERF9). AGI codes of mTERF9 protein interactors can be found in Supplementary Data Set 1. The T-DNA mutant used was WiscDsLox474E07 (*mterf9*).

## RESULTS

### mTERF9 is a chloroplast nucleoid-associated protein required for plant growth

To characterize the molecular function of mTERF9, we analyzed the *Arabidopsis mterf9* mutant that was previously reported to be affected in plant development (24). This mutant carries a T-DNA insertion in the fourth intron of the *mTERF9*/At5g55580 gene (Figure 1A). *mTERF9* encodes a 496 amino acid protein harboring seven tandem mTERF motifs that are preceded by a predicted N-terminal chloroplast transit peptide (Figure 1A). We confirmed the *mterf9* mutant phenotype at different developmental stages. *mterf9* plants exhibited a pale leaf pigmentation and a slower growth phenotype compared to wild-type (WT), but remained fertile (Figure 1B). The introduction of a WT copy of the *mTERF9* gene under the control of the CaMV 35S promoter into *mterf9* fully restored the WT phenotype demonstrating that the mutant phenotype resulted from *mTERF9* disruption. RT-PCR analysis confirmed the lack of *mTERF9* full-length mRNA in *mterf9* and its restoration in the complemented *mterf9* plants (CP) (Figure 1C). The chlorotic phenotype displayed by *mterf9* suggests a potential loss of photosynthetic activity in the mutant. Therefore, the functional status of photosynthesis of the mutant was monitored using a pulse amplitude modulated system (Table 1). In all respects, the complemented lines showed characteristics comparable to the WT. The *mterf9* mutant displayed a decrease in photosystem II (PSII) activity as revealed by a reduced maximum quantum yield of PSII (0,70 vs 0,81; *mterf9* vs WT) and an increased minimum fluorescence value (Fo) (Table 1). Effective quantum yield of PSII measured in the steady state 5 min after induction was decreased from 0,73 in the WT to 0,58 in *mterf9* whereas non-photochemical quenching was not affected. Overall, photosystem I (PSI) activity was reduced by one third in mutant plants as compared to the WT and no PSI donor side limitation could be detected. Instead, the quantum yield of non-photochemical energy dissipation due to PSI acceptor side limitation was reduced by about a half. The data indicate and confirmed a pleiotropic photosynthetic deficiency in the *mterf9* mutant rather than a specific defect. To confirm the predicted chloroplast intracellular localization of mTERF9, we transiently expressed an mTERF9 protein fused to a C-terminal GFP in *Nicotiana benthamiana* leaves and examined leaf protoplasts by confocal microscopy (Figure 1D). The results revealed that the fusion protein localizes to punctuated foci overlapping with the chloroplast chlorophyll autofluorescence and additionally, with the fluorescence of a co-expressed nucleoid-associated chloroplast protein, RAP fused with RFP (39). These results indicate that mTERF9 functions in chloroplasts of plant cells where it associates with the nucleoid.

**Figure 1.**
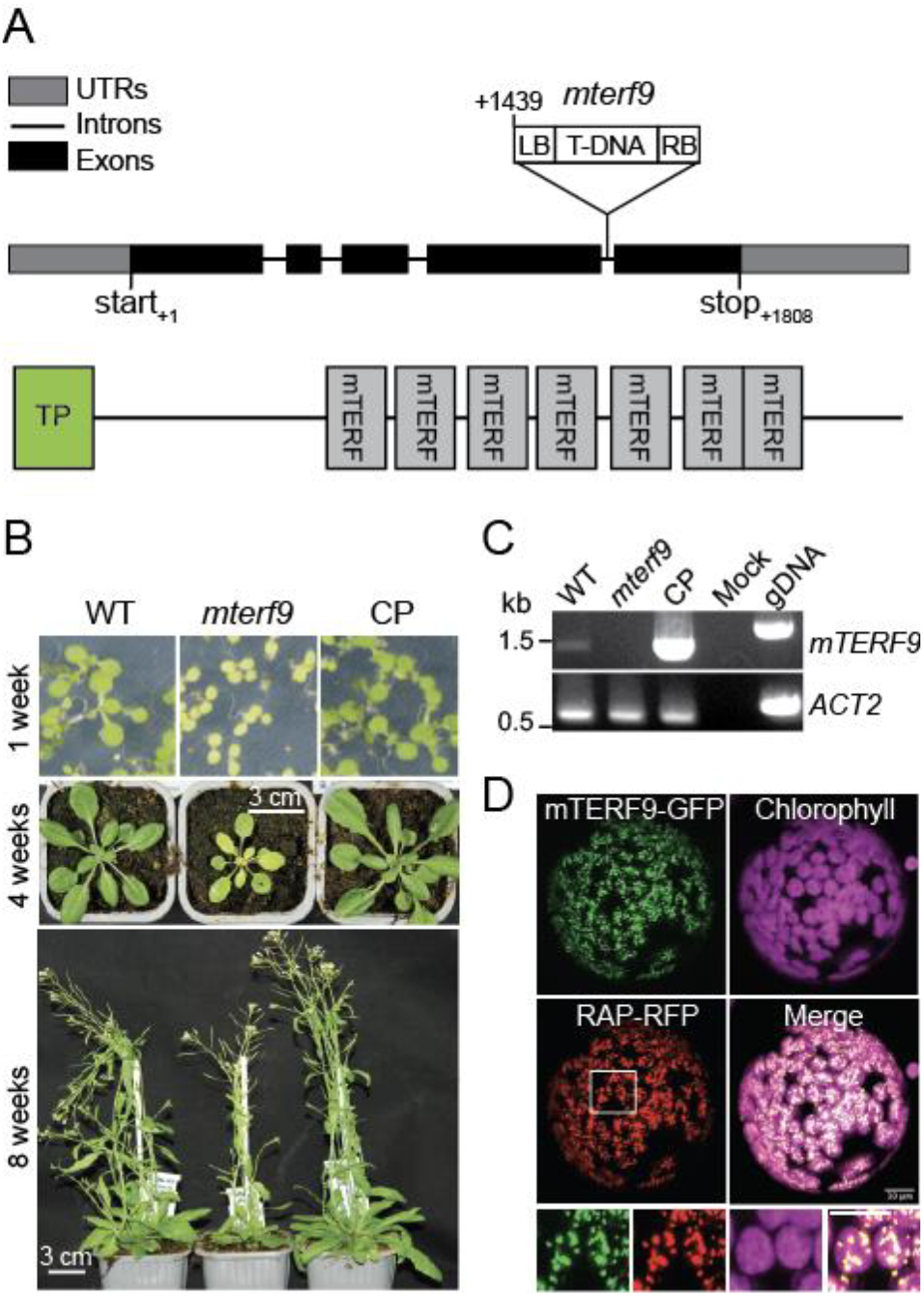
mTERF9 is a chloroplast nucleoid-associated protein required for plant development. **(A)** Schematic representation of the *mTERF9* gene and protein with the position of the *mterf9* T-DNA insertion. **(B)** Phenotypes of wild-type (WT), *mterf9* and complemented (CP) plants grown in medium or soil at indicated growth stages. **(C)** RT-PCR analysis of *mTERF9* expression in WT, *mterf9* and complemented plants. Genomic DNA (gDNA) was used as positive control for PCR and *ACTIN-2* (*ACT2*) serves as internal control for RT-PCR. **(D)** Subcellular localization of mTERF9-GFP and RAP-GFP fusion proteins in tobacco leaf protoplasts. Close-up views of the framed area are shown below. Scale bar: 5 μm.

**Table 1.** Chlorophyll *a* fluorescence induction and light-induced PSI absorbance changes. Chlorophyll *a* fluorescence was measured on 2-week-old Arabidopsis plants grown on soil. Representative measurements of chlorophyll *a* fluorescence in the WT, *mterf9* mutants and complemented (CP) lines. Saturating light pulses were given in 20 s intervals during induction (27).

|  | WT<sup>h</sup> (n<sup>i</sup>=5) | *mterf9* (n=5) | CP (n=5) |
| --- | --- | --- | --- |
| **Fv/Fm<sup>a</sup>** | 0,81 ± 0,00 | 0,70 ± 0,01 | 0,81 ± 0,01 |
| **Fo** | 1,00 ± 0,11 | 1,25 ± 0,05 | 1,02 ± 0,08 |
| **Φ<sub>PSII</sub><sup>b</sup>** | 0,73 ± 0,01 | 0,58 ± 0,01 | 0,72 ± 0,02 |
| **NPQ<sup>c</sup>** | 0,20 ± 0,02 | 0,22 ± 0,02 | 0,20 ± 0,02 |
| **Φ<sub>PSI</sub><sup>d</sup>** | 0,47 ± 0,02 | 0,42 ± 0,05 | 0,41 ± 0,01 |
| **Φ<sub>PSI ND</sub><sup>e</sup>** | 0,27 ± 0,05 | 0,48 ± 0,03 | 0,29 ± 0,04 |
| **Φ<sub>PSI NA</sub><sup>f</sup>** | 0,28 ± 0,03 | 0,12 ± 0,05 | 0,30 ± 0,03 |
| **%<sup>g</sup> ΔA<sub>P700</sub>** | 100,00 ± 11,22 | 67,34 ± 4,65 | 104,51 ± 14,5 |

<sup>a</sup> maximum quantum yield of PSII
<sup>b</sup> effective quantum yield of PSII (50 μmol photons m<sup>-2</sup> s<sup>-1</sup>).
<sup>c</sup> non-photochemical quenching.
<sup>d</sup> quantum yield of PSI.
<sup>e</sup> quantum yield of non-photochemical energy dissipation due to PSI donor side limitation.
<sup>f</sup> quantum yield of non-photochemical energy dissipation due to PSI acceptor side limitation.
<sup>g</sup> maximum absorbance of P700 in % of the WT.
<sup>h</sup> the data for the WT were collected along with *mterf9* and an additional Arabidopsis mutant *mda1* (21). As such, the WT values are the same as in (21).
<sup>i</sup> number of plants measured.

### mTERF9 deficiency impairs chloroplast protein accumulation and translation

The pale leaf and defective photosynthesis phenotypes displayed by the *mterf9* mutant suggests an mTERF9 function related to chloroplast biogenesis. To investigate mTERF9 function in chloroplasts, we first analyzed the accumulation of representative subunits of chloroplast protein complexes by immunoblotting in the mutant. With the exception of the plastid-encoded RpoB and nuclear-encoded LHCB2, FBA, CPN60α1 and β1 proteins, the results showed a ~50-75% decrease in the amount of chloroplast proteins tested in *mterf9* compared to WT and CP plants (Figure 2A). We additionally confirmed the expression of the mTERF9 protein fused to a C-terminal 4xMyc tag in the CP plants and showed its dual-detection in both the stroma and membrane fractions of chloroplasts (Figure 2A and 2B). The global reduction of the amount of chloroplast protein complexes in *mterf9* including the plastid ribosomal protein S1 (RPS1) suggests a possible defect in chloroplast translation and ribosome biogenesis. To confirm this, we investigated the *de novo* synthesis of chloroplast proteins by protein pulse-labeling with ^35^S-methionine. The results showed that the synthesis rates of RbcL and D1 proteins were lower in *mterf9* with a respective ~25 and 80% decrease relative to WT and CP plants, respectively (Figure 2C). Overall, the results indicated that the loss of mTERF9 activity impairs the accumulation of chloroplast proteins and translation.

**Figure 2.**
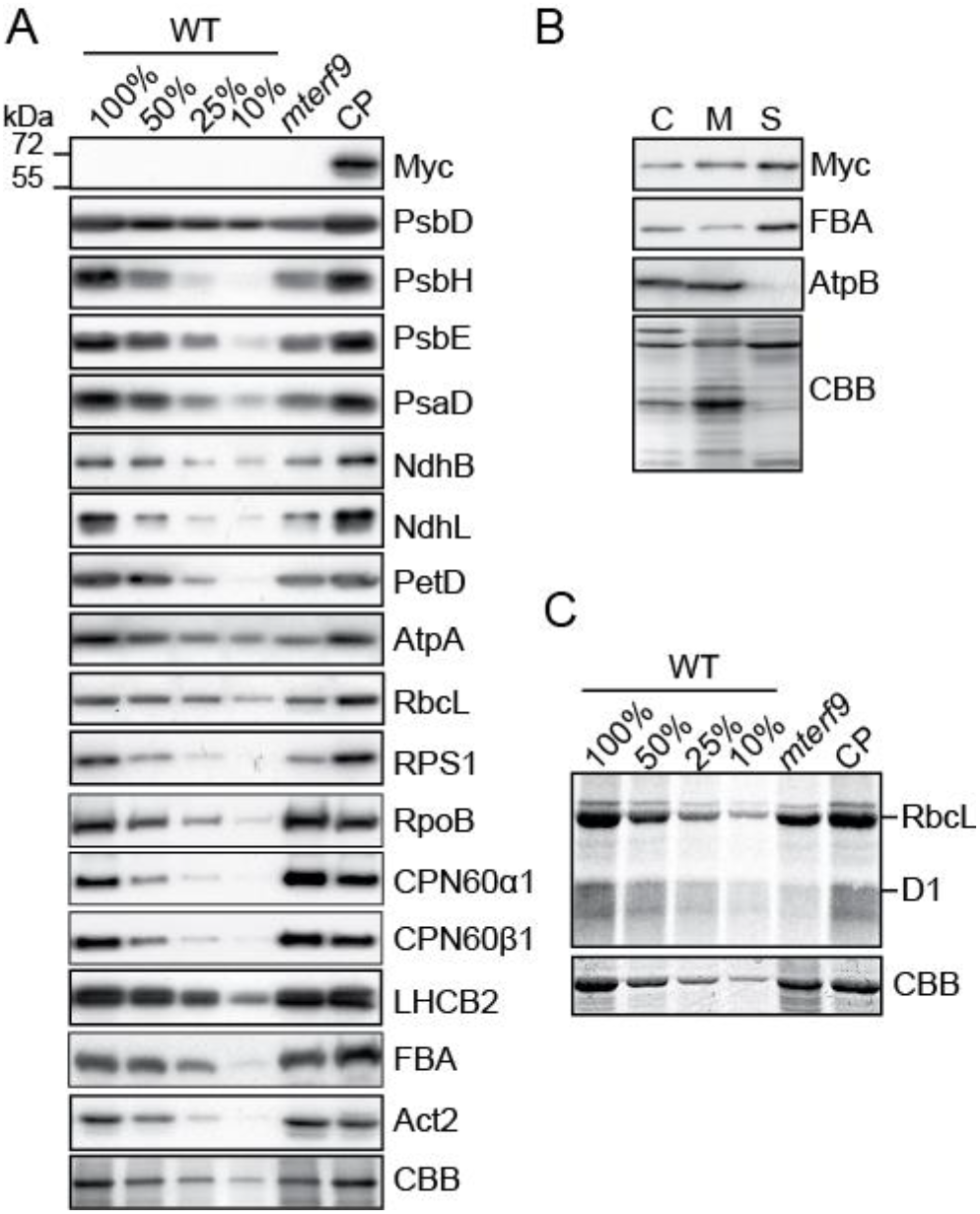
Chloroplast protein accumulation deficiency in *mterf9*. **(A)** Immunoblot analyses of total leaf protein extracts with antibodies against mTERF9-Myc and subunits of the photosystem I (PsaD), photosystem II (PsbD, PsbH, PsbE,), Cytochrome *b*_*6*_*f* (PetD), NADH dehydrogenase (NdhB, NdhL), ATP synthase (AtpA), Rubisco (RbcL), light-harvesting complex II (LHCB2), chloroplast ribosome (RPS1), Plastid-encoded RNA polymerase (RpoB), chaperonin 60 (CPN60α1, CPN60β1) complexes and the stromal fructose-bisphosphate aldolase 1 enzyme (FBA). Replicate membranes were stained with Coomassie Blue (CBB) to show equal protein loading. The PsbD, PsbH, PsbE, PsaD, NdhB, NdhL, PetD, AtpA, RbcL, RPS1 immunoblot images for the WT were previously reported (21) and are reproduced here with permission. **(B)** mTERF9 localizes to the stroma and chloroplast membranes. Isolated chloroplasts were lysed in hypotonic buffer and membrane and soluble protein fractions were separated by centrifugation. Chloroplast (C), soluble (S), and membrane (M) protein fractions were analyzed by immunoblotting using antibodies against Myc epitope, a stromal protein (FBA) and a membrane associated subunit of the ATP synthase complex (AtpB). The Coomassie Blue (CBB) stained membrane is shown. **(C)** *In vivo* chloroplast translation assays. Leaf discs from the indicated genotypes were pulsed-labelled with ^35^S-Methionine and neosynthesized proteins were separated by SDS-PAGE and visualized by autoradiography. The CBB stained gel is shown below and serves as loading control.

### mTERF9 deficiency causes reduced accumulation of the *16S* and *23S* rRNAs

Members of the mTERF family are predicted to control gene expression in organelles (6,9) and the loss of chloroplast translational activity in *mterf9* can result from the altered expression of some chloroplast genes. To identify which genes were affected in *mterf9*, we measured chloroplast gene transcripts by qRT-PCR (Figure 3) and found that the steady-state levels of mRNAs were moderately increased (0<log_2_FC<2.5) or unchanged in the *mterf9* mutant. The transcript overaccumulation in *mterf9* was confirmed by northern blot analysis for selected genes using complementary probes against *matK*, *ndhD*, *rbcL*, *ycf3* and *rpoC1* transcripts (Supplementary Figure 1). In contrast, the *16S* and *23S* rRNAs, two RNA constituents of the chloroplast small and large ribosomal subunits were reduced compared to the WT and CP plants as observed by qRT-PCR (Figure 3). The rRNAs are unstable when not incorporated into the chloroplast ribosomal subunits and therefore, their reduction in *mterf9* is indicative of a partial loss of chloroplast ribosomes content in the mutant. A global increase in the steady-state levels of chloroplast mRNAs has previously been reported in plants whose chloroplast translation is chemically or constitutively impaired (40,41). Therefore, the moderate increase of chloroplast transcripts in *mterf9* is likely a secondary effect of reduced chloroplast translation. Some mTERF proteins have been involved in RNA intron splicing in plant organelles and a lack of splicing for some chloroplast genes can lead to translation impairment when these encode components that are important for the ribosome biogenesis. Thus, we additionally assayed the intron splicing efficiency for chloroplast genes in *mterf9* relative to WT by qRT-PCR (Supplementary Figure 2). At the exception of a slight reduction for *ycf3* intron 1, splicing was not significantly disrupted in *mterf9*. However, northern blot analyses using *ycf3* strand specific probes were not consistent with the qRT-PCR results and showed very little, if any splicing defect in *ycf3* intron 1 (Supplementary Figure 1). Instead, the northern blot results showed an overexpression of *ycf3* pre-mRNAs as discussed previously. Therefore, neither the transcripts over-accumulation nor intron splicing defects in *mterf9* can explain the overall reduced accumulation of chloroplast proteins and translation in *mterf9*. By contrast, the observed decrease of the *16S* and *23S* rRNAs in *mterf9* indicate that mTERF9 is required for the accumulation of chloroplast ribosomes, which is congruent with the global reduction of chloroplast-encoded proteins in *mterf9*.

**Figure 3.**
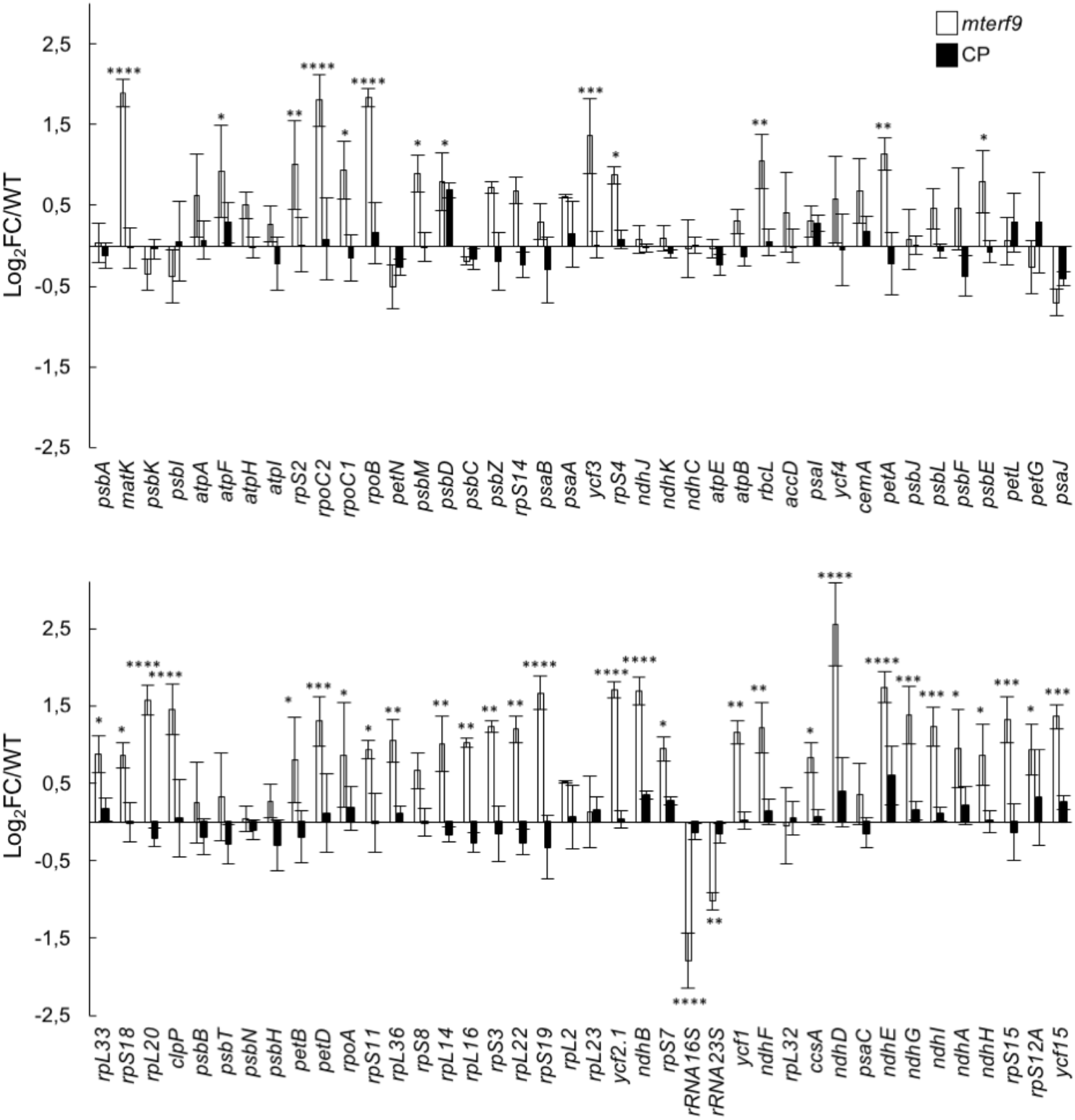
Steady state levels of chloroplast gene transcripts in Arabidopsis in *mterf9* and CP plants. Transcript levels were determined by qRT-PCR and are displayed as the log_2_ fold change (FC) relative to WT for the mutant or the CP plants. Genes are ordered according to their genome positions. The nuclear *ACT2* and *TIP41* genes were used for data normalization. The values from three biological replicates performed each with technical triplicate were averaged per genotype and standard errors are indicated. ANOVA Dunnet’s multiple test correction: *, *P* < 0.05; **, *P* < 0.005; ***, *P* < 0.0005; ****; *P* < 0.00005.

### mTERF9 is required for the accumulation of *16S* and *23S* rRNAs

Chloroplast rRNA genes are organized in an operon and the *16S* and *23S* rRNAs are co-transcribed with the *4.5S* and *5S* rRNAs leading to RNA precursors that are subjected to a series of processing events (reviewed in 42) (Figure 4). For example, the *23S* rRNA is internally fragmented at two “hidden breaks”, leading to the accumulation of seven distinct transcripts (Figure 4A). To further investigate and confirm the decrease of the *16S* and *23S* rRNAs in *mterf9*, RNA gel blot analyses were conducted in biological triplicates and signals were quantified (Figure 4 and Supplementary Figure 3) with probes designed to detect each rRNA and their processed forms (Figure 4A). The results confirmed the ~50% reduction in the abundance of the processed 1.5 kb *16S* rRNA in *mterf9* compared to the WT or complemented plants (Figure 4B and 4C). RNA gel blot hybridization with three probes designed to detect the different fragments of the 23S rRNA revealed significant reduction of the 2.4, 1.3, 1.1 and 0.5 kb *23S* rRNAs in *mterf9* with a pronounced effect on the 2.4 kb isoform (~60% reduction). In addition, the accumulation of the processed *4.5S* and *5S* rRNAs were not significantly affected in the mutant. The RNA gel blotting results confirmed the rRNA deficiencies in *mterf9* and the importance of mTERF9 for the accumulation of the *16S* and *23S* rRNAs *in vivo*.

**Figure 4.**
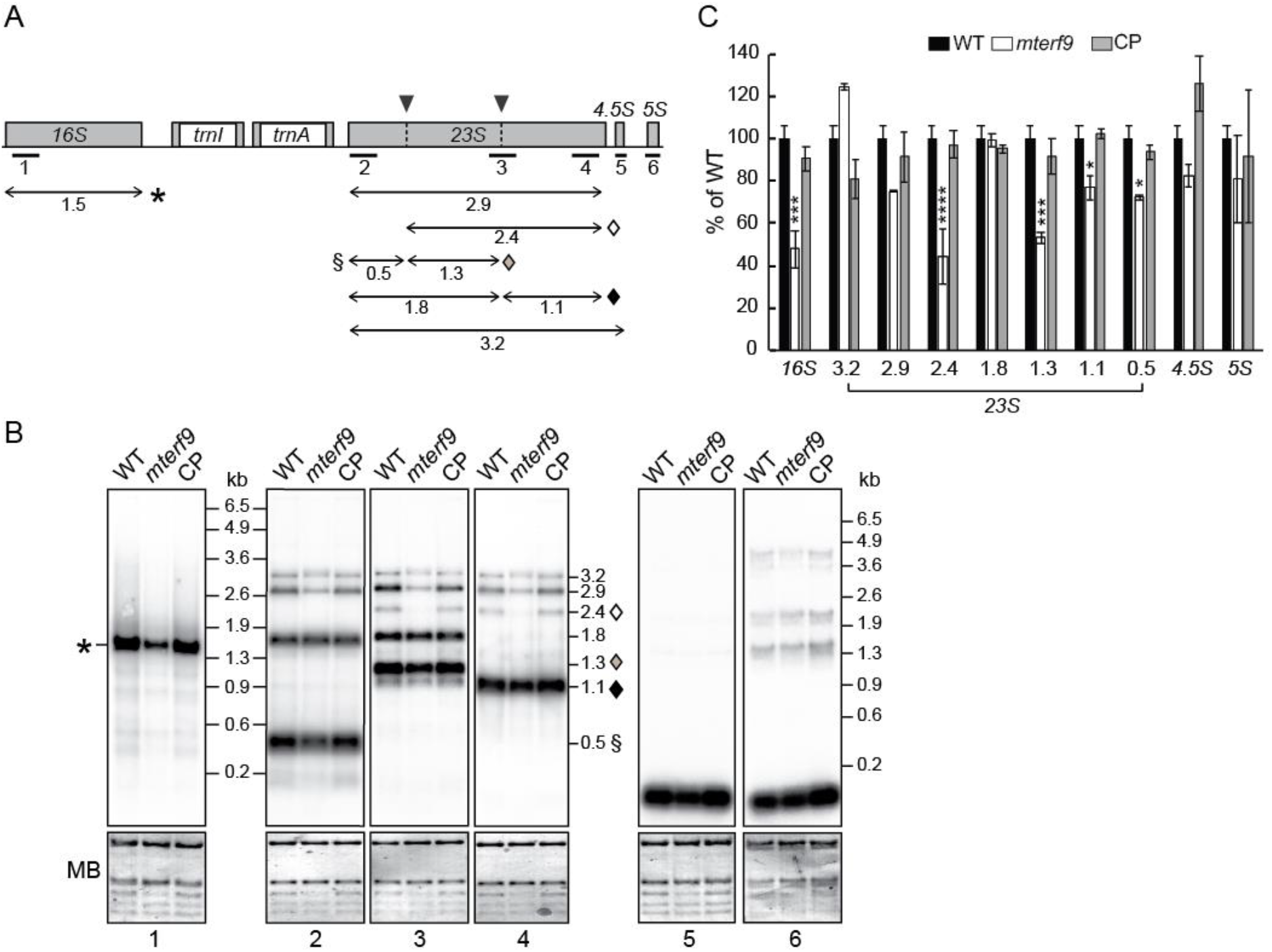
Defects in the *16S* and *23S* rRNA accumulation in *mterf9* plants. **(A)** Schematic representation of the chloroplast rRNA operon. Exons and introns are represented by gray and white boxes, respectively. The positions of the probes used for RNA blot hybridization are indicated beneath the map. The *23S* rRNA hidden breaks positions are shown above the gene with black arrowheads. The major accumulating transcripts for the *16S* and *23S* rRNA genes are mapped with arrows and their size is given in kb below. Transcripts specifically impaired in the *mterf9* are indicated with symbols. **(B)** Total leaf RNA (0.5 μg) was analyzed by RNA gel blot hybridization using probes diagrammed in (A). An excerpt of the RNA membranes stained with methylene blue (MB) are shown to illustrate equal RNA loading, respectively. **(C)** Relative quantification to WT in the accumulation of chloroplast rRNAs in *mterf9* and CP plants. The average and standard error of three biological replicates is shown. Fisher’s test: *, *P* < 0.05; ***, *P* < 0.0005; ****, *P* < 0.00005.

### mTERF9 associates with the ribosomal 30S subunit to promote ribosomal assembly and translation

The deficiency in the rRNAs accumulation in *mterf9* points towards a reduction in the chloroplast ribosome content in the mutant and a possible defect in ribosomal assembly. The chloroplast 70S ribosome is composed of the small 30S and large 50S subunits that respectively contain the *16S* and the *23S*, *4.5S* and *5S* rRNAs. Preliminary immunoblotting analysis indicated a partial loss of RPS1, a protein of the 30S subunit (Figure 2A). We analyzed the sedimentation of the 30S and 50S ribosome subunits in *mterf9*, WT and CP plants by sucrose gradient sedimentation of stromal protein complexes (Figure 5A). The fractionation of the 30S and 50S ribosomal subunits on the gradient were monitored by immunoblotting using antibodies against RPS1, RPS7 and RPL33. In the WT and CP plants, RPL33 mostly sedimented in the last fractions of the gradient (fractions 10 to pellet), whereas RPS1 and RPS7 sedimented in the middle of the gradient (peak fractions 5 to 7 and 7 to 10, respectively). By contrast, in *mterf9*, RPL33, RPS1 and RPS7 sedimentation patterns were shifted to lower molecular-weight fractions, with a more pronounced shifting for RPS1. These results demonstrate that the loss of mTERF9 function in Arabidopsis compromised the assembly of the chloroplast ribosome. Additional immunoblot analysis with an antibody against the Myc tag showed that mTERF9 co-sedimented predominantly with RPS1 and RPS7, indicating that it is found in particles of the same size than the 30S ribosomal subunit in chloroplasts.

**Figure 5.**
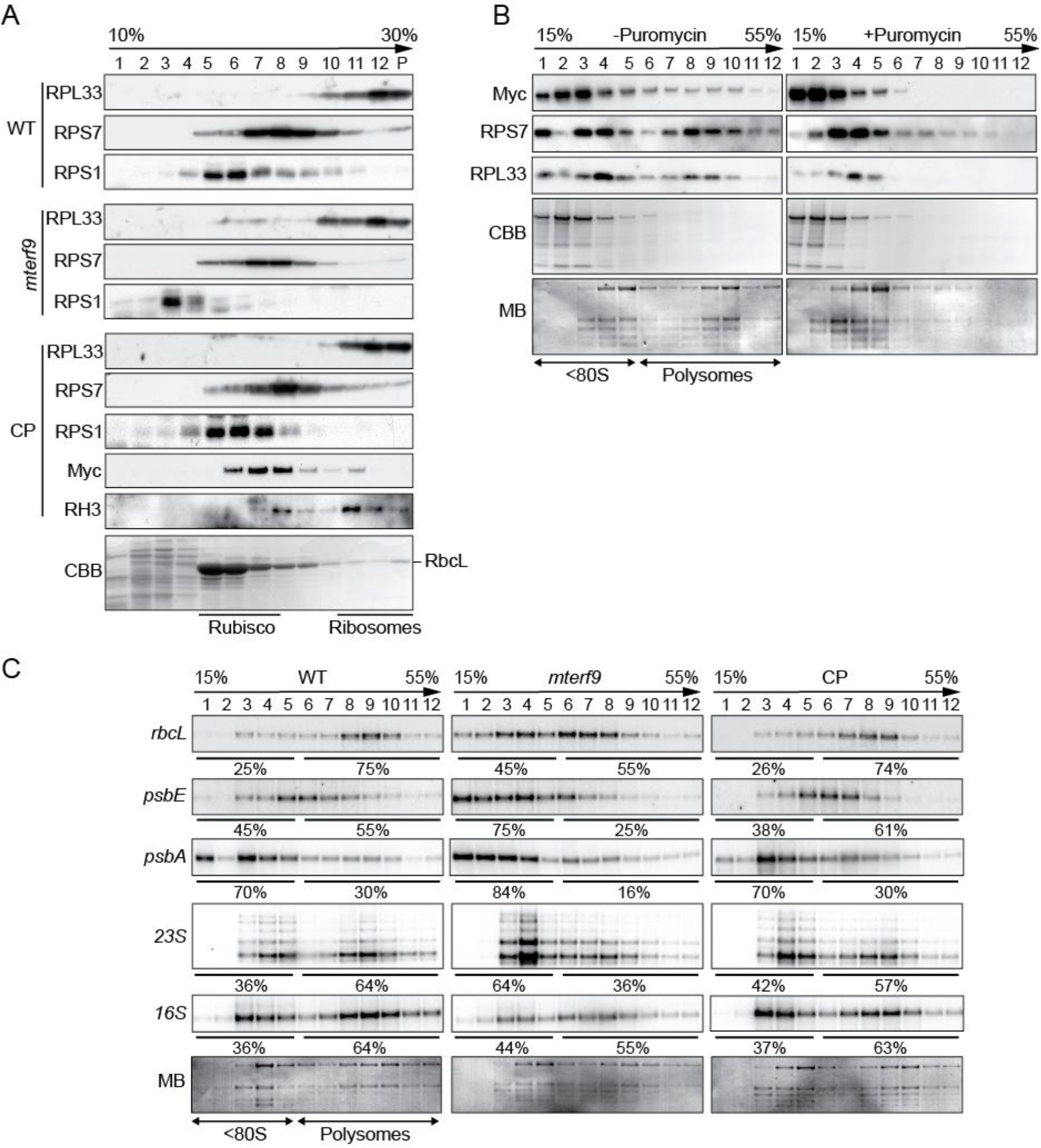
mTERF9 associates with chloroplast ribosomes and promotes mRNA association with ribosomes *in vivo*. **(A)** Sucrose gradient fractionation of stroma from the indicated genotypes. An equal volume of each fraction was analyzed by immunoblots with antibodies against ribosomal proteins from the small (RPS1, RPS7) and large subunit (RPL33), mTERF9 (Myc). The Coomassie blue-stained membrane (CBB) of the WT fractions is shown below. **(B)** Polysomal association of mTERF9. Leaf polysomes from complemented *mterf9* plants were fractionated on 15% to 55% sucrose gradients. Fractions were analyzed by immunoblotting with antibodies against mTERF9 (Myc) and ribosomal proteins (RPS7 and RPL33) and RNA electrophoresis on denaturing agarose gel (bottom). The protein and RNA membranes stained with CBB and MB are shown, respectively. The sedimentation of the <80S ribosomes and polysomes on the gradient was confirmed with a puromycin control. **(C)** Polysome loading of selected chloroplast RNAs in WT, *mterf9* and CP plants. Fractions from sucrose density gradients were analyzed by RNA gel blots using gene-specific probes. Signals in the polysomal (fractions 6-12) and monosomes/free RNA fractions (fractions 1-5) were quantified by phosphor-imaging and are displayed as percentage of the total signal over the 12 fractions.

We next determined whether mTERF9 associated with chloroplast ribosomes engaged in translation by polysome analysis from sucrose gradients (Figure 5B). The polysome-containing fractions were identified by immunodetection with the RPS7 and RPL33 antibodies and by visualization of the cytosolic rRNAs by RNA electrophoresis on a denaturing agarose gel. The polysomes were detected in fractions 6 to 12 and mTERF9 was detected in these fractions as well as in fractions containing monosomes and immature ribosomal particles (fractions 1 to 5). Treatment of the polysomes with the dissociating agent puromycin prior to their fractionation on sucrose gradient efficiently released mTERF9 from heavy to lighter complexes containing mostly monosomes or immature ribosomal particles, confirming the association of mTERF9 with the polysomes.

Finally, we analyzed the association of chloroplast mRNAs and the *16S* and *23S* rRNAs with polysomes in the WT, *mterf9* and CP plants by sucrose density gradient fractionation and northern blot analyses (Figure 5C). As shown by the levels of mature *16S* and *23S* in polysomal fractions (fractions 6 to 12), *mterf9* contained fewer polysomes than the WT and CP plants. In addition, the RNA gel blot results showed that the *rbcL*, *psbE* and *psbA* transcripts were partially shifted to the top of the gradient in *mterf9* compared to WT, indicating that their loading to the polysomes and their translation efficiency were diminished in the mutant. These results correlate well with the lower rate of chloroplast protein synthesis that was observed in *mterf9* (Figure 2C).

Altogether, the results show that mTERF9 is required for chloroplast ribosomal assembly and translation. mTERF9 primarily associates with the 30S subunit that assembles with the 50S to form the functional 70S chloroplast ribosome. In addition, mTERF9 association with the polysomes indicates that the protein plays a role during translation.

### mTERF9 binds the *16S* rRNA *in vivo*

mTERF proteins are nucleic acid binding proteins that have been predominately involved in DNA-related functions in organelles. However, this paradigm has recently shifted with the reports of two mTERF proteins involved in RNA intron splicing in plant organelles (43,44). The chloroplast rRNA defects in *mterf9* and mTERF9 co-sedimentation with the ribosomes suggest that the protein target the rRNAs *in vivo*. To explore this possibility and identify mTERF9 RNA ligands *in vivo*, we performed genome-wide RNA co-immunoprecipitation assays (RIP). Stromal extracts from the CP and WT plants were subjected to immunoprecipitation with antibodies raised against the Myc tag and co-immunoprecipitated RNAs were identified by hybridization to tiling microarrays of the Arabidopsis chloroplast genome (RIP-chip) (Figure 6A). The results revealed a prominently enriched peak (>20-fold) in the mTERF9 immunoprecipitate that corresponds to the *16S* rRNA and minor peaks (<10-fold) in the *23S*, *4.5S* and *5S* rRNAs as well as *atpH*, *psbC* and *psbE* loci. To quantify mTERF9 binding to these RNA targets, we conducted an independent RIP experiment followed by qRT-PCR analysis of the immunoprecipitated RNAs (Figure 6B). The results confirmed that mTERF9 significantly binds to the *16S* rRNAs and to a lesser extent the *23S* rRNA. However, *atpH*, *psbC* and *psbE* were not significantly enriched in mTERF9 immunoprecipitate as compared to two negative control genes, *rpoB* or *rbcL* indicating that these targets were either false positives or unstable ligands (Figure 6B). Taken together, the results confirmed that mTERF9 primarily binds the *16S* rRNA *in vivo*, which is consistent with its association with the small 30S ribosomal subunit.

**Figure 6.**
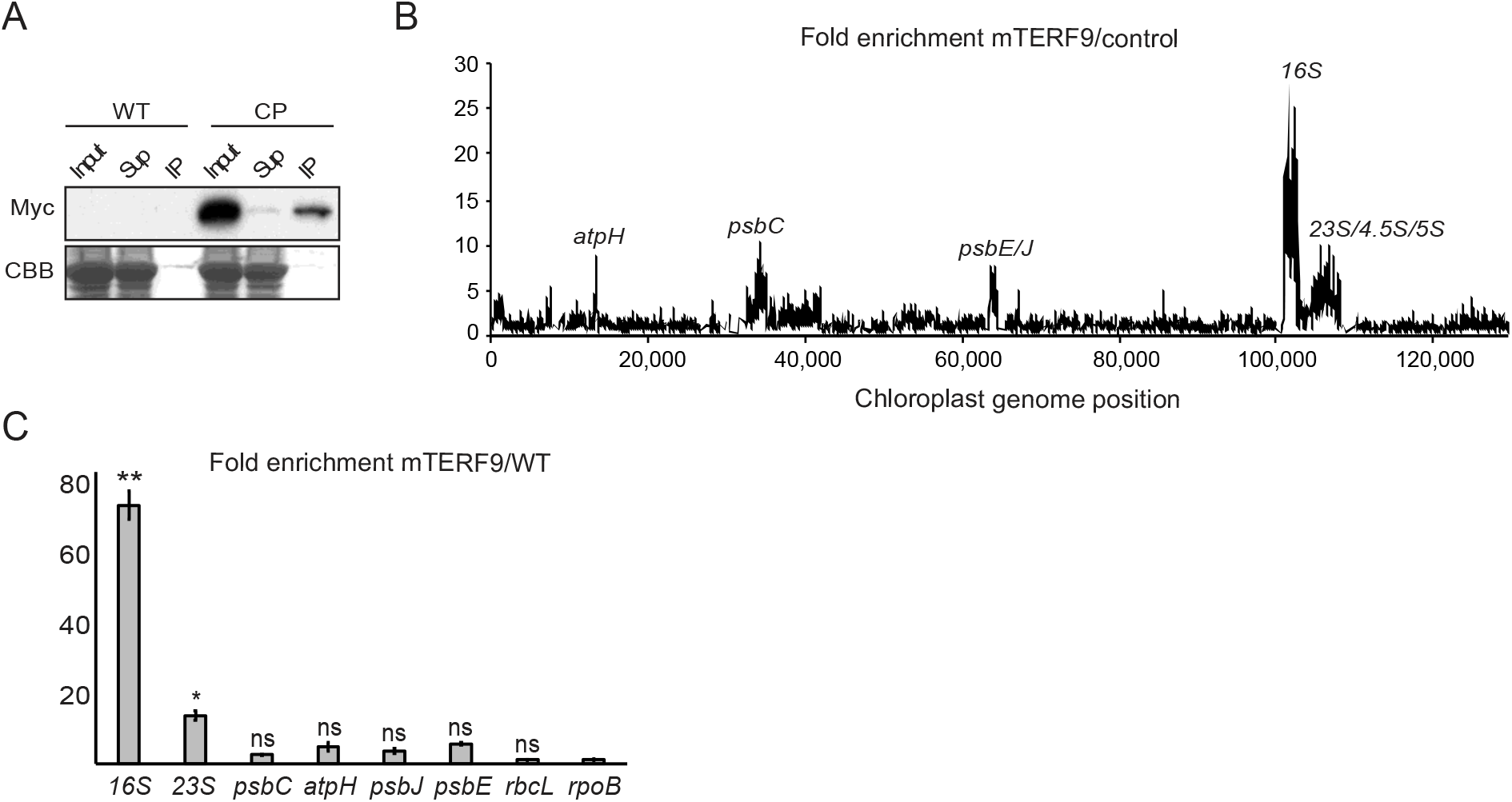
mTERF9 associates with the *16S* rRNA in chloroplasts. **(A)** mTERF9 RNA ligands were identified by co-immunoprecipitation on stromal extract from the complemented *mterf9* (CP) or wild-type (WT, negative control) with anti-Myc antibodies, followed by RNA hybridization on a chloroplast genome tiling microarray. The efficiency of mTERF9 immunoprecipitation was confirmed by immunoblot analysis with anti-Myc antibodies (left panel). Sup: supernatant, IP: immunoprecipitate. The enrichment ratios (ratio of signal in the immunoprecipitation pellet versus supernatant) are plotted according to position on the chloroplast genome after subtracting values obtained in the negative control immunoprecipitation (WT stroma) (right panel). The RIP-chip assay revealed the predominant enrichment of the *16S* rRNA in mTEF9 immunoprecipitate. **(B)** Validation of mTERF9 RNA *in vivo* ligands by qRT-PCR. The levels of immunoprecipitated RNAs were calculated as percent recovery of the total input RNA in control WT IP and mTERF9 IP in triplicate, and the average ratio and standard error are shown. Kruskal-Wallis test; * = *P* < 0.05.

### The mTERF9 protein interactome confirms its link to ribosome biogenesis

Our data demonstrated that mTERF9 is involved in ribosomal assembly and chloroplast translation. The recruitment of components of the ribosome biogenesis machinery to the *16S* rRNA may be one of the key functions of an rRNA-interacting protein. To understand the protein interactome of mTERF9 *in vivo* and confirm its *in vivo* association with the chloroplast ribosome, we performed co-immunoprecipitation of untreated or RNase-treated stromal extracts in biological triplicates and, proteins from the immunoprecipitated fractions were identified by LC-MS/MS. The efficiency of mTERF9-Myc immunoprecipitation between the RNase-treated or untreated samples was similar, allowing a direct comparison of the results (Figure 7A). We identified 158 and 173 proteins significantly enriched by mTERF9-Myc precipitation (Log_2_(FC)>2 and adj_*P*<0.05) in the −RNase and +RNase condition, respectively (Figure 7B and 7C). The enriched mTERF9-interacting proteins were classified in 7 groups according to their functional annotations: ribosomal proteins of the small and large subunits (RPSs and RPLs), CPN60 chaperonins, rRNA processing/ translation factors, RNA binding proteins (RBPs), components of the transcriptional active chromosome (TACs) and finally, the category “others” grouping chloroplast proteins with functions unrelated to gene expression and cytosolic protein contaminants (Figure 7C; Supplementary Data Set 1). Gene ontology term enrichment analyses revealed that the RNA-dependent and -independent protein interactants share over-represented molecular functions in ribosome biogenesis (Figure 7D). As an illustration, the *in vivo* fishing of mTERF9 in the absence of RNase treatment pulled down 20 out of the 24 proteins that constitute the small ribosome subunit and 27 out of the 33 ribosomal proteins composing the large subunit of chloroplasts (45). Nevertheless, the RNase treatment had differential effects on the accumulation of proteins in mTERF9 co-immunoprecipitates. The treatment reduced the number of chloroplast ribosomal proteins and in particular of the large subunit, rRNA processing/translation factors and TAC components, while it increased the number of RNA binding proteins and proteins from the category “others” (Figure 7C). On contrary, all 6 subunits (CPN60α1-2, CPN60β1-4) of the chloroplast CPN60 chaperonin complex (46) were constantly retrieved in both conditions as most enriched proteins in mTERF9 co-immunoprecipitates (Figure 7B and 7E). In total, 92 proteins were commonly found in the untreated and RNase-treated co-immunoprecipitates (Figure 7E) suggesting that the majority of the mTERF9 interactants were not RNA-dependent but rather direct protein interactors. However, this does not exclude the possibility that the remaining co-immunoprecipitated proteins engaged in direct protein-protein interactions with mTERF9. Among the 92 common proteins, none were chloroplast RNA binding proteins, indicating that the RNase treatment efficiently destabilized ribonucleoprotein complexes and that their interaction with mTERF9 was RNA-dependent. Twelve proteins from the small and large ribosomal subunits were respectively enriched under both conditions along with 6 chloroplast rRNA processing and translation factors (Figure 7E and Supplementary Data Set 1). These include the following rRNA processing factors: RNA helicases, RH3 (47,48) and ISE2 (49), Ribonuclease J RNJ (50), RNA binding protein RHON1 (51) and the translation initiation and elongation factors FUG1 (52) and EF-Tu/SVR11 (53), respectively. Finally, ten TAC components co-immunoprecipitated with mTERF9 under both conditions. The TACs enrichment in mTERF9 co-immunoprecipitates was consistent with their co-localization to the nucleoids, a site known to play a major function in rRNA processing and ribosome assembly in chloroplasts (39,54,55). Interestingly, some ribosomal proteins, rRNA processing factors and RNA binding proteins were exclusively co-immunoprecipitated with mTERF9 by RNase treatment (Figure 7E and Supplementary Data Set 1). The RNase-dependency of these interactors revealed that their interaction with mTERF9 occurred upon mTERF9 dissociation from ribosomal nucleoprotein complexes which suggests that mTERF9 can interact with ribosomal proteins and rRNA processing factors in a spatial and sequential order during the assembly/disassembly of the ribosome subunits *in vivo*.

**Figure 7.**
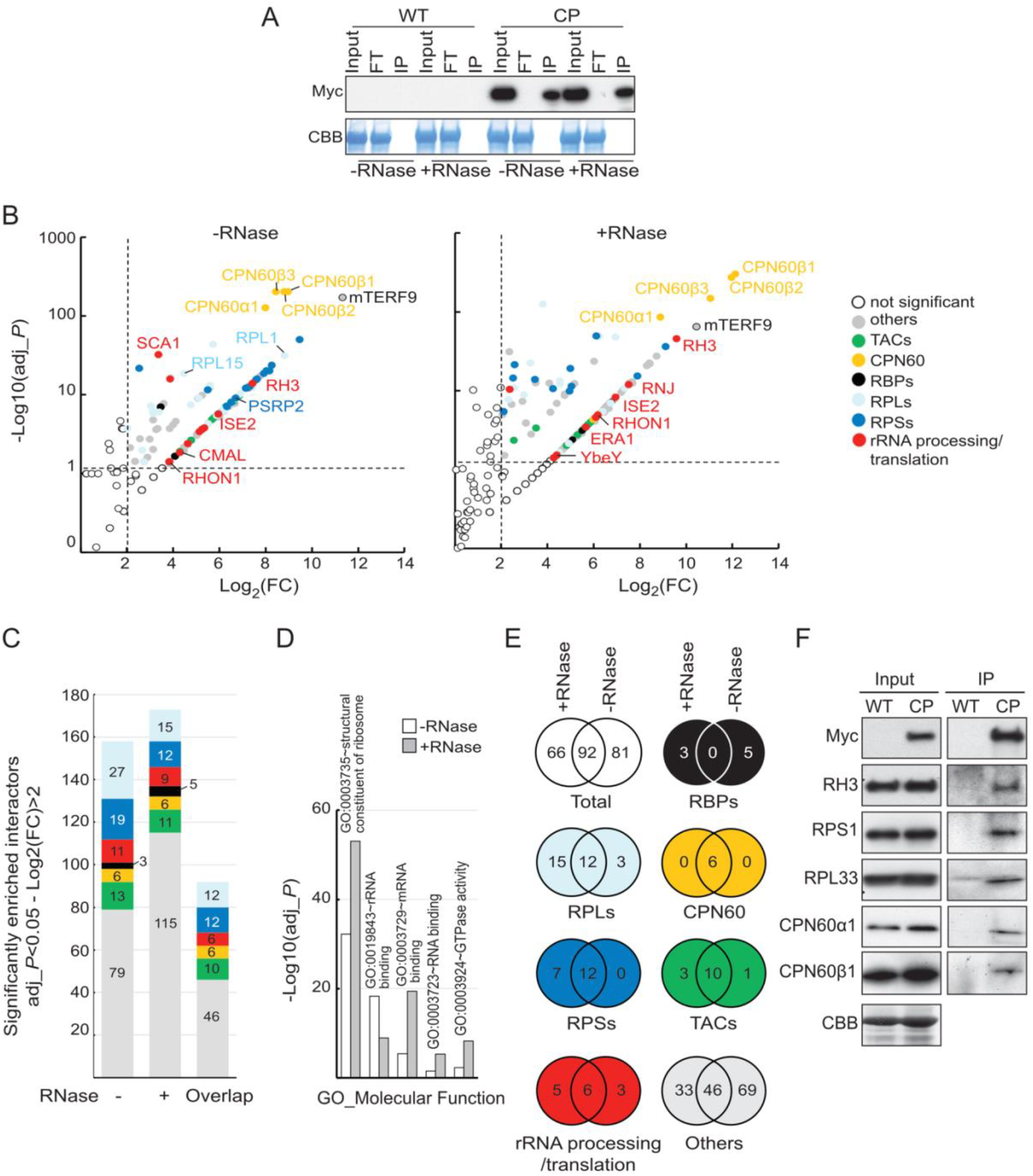
mTERF9 protein interactome is highly enriched with proteins involved in chloroplast ribosome biogenesis. **(A)** mTERF9 immunoprecipitation. Untreated or RNase-treated stroma extracts from complemented *mterf9* (CP) or wild-type (WT) plants were used for immunoprecipitation with anti-Myc antibody. The input, flow-through (FT) and immunoprecipitate (IP) fractions were analyzed by immunoblot with anti-Myc antibody. A portion of the Coomassie blue-stained membrane (CBB) showing the abundance of RbcL as loading control. **(B)** Volcano plots show the enrichment of proteins co-purified with mTERF9 and identified by mass spectrometry in absence or presence of RNase in comparison with control IPs. IPs were performed on biological triplicate. *Y*- and *X-*axis display Log10 scale of −Log_10_ adjusted *p*-values (adj*_P*) and Log_2_ fold changes (FC) of proteins, respectively. The dashed lines indicate the threshold above which proteins were significantly enriched (*p*-value <0.05 and FC >4). Proteins are color-shaded according to their functional group and the color key provided to the right. ns: not significant. The full lists of mTERF9-associated proteins and their Arabidopsis locus identifiers are available in Supplementary Data Set 1. **(C)** Bar chart showing the number of significant mTERF9 interacting proteins in the functional groups. The same color code than in (B) is used. The “overlap" bar represents common proteins found in mTERF9 protein interactomes in absence of presence of RNase. **(D)** Bar chart depicting the functional analysis of the mTERF9 protein interactomes and showing the 5 terms contained in the top functional annotation cluster identified by DAVID gene analysis online tool using the default parameters (88). GO terms are plotted according to −Log_10_ of their respective adjusted *p*-values. **(E)** Venn diagrams showing the significantly enriched proteins in each functional category in mTERF9 immunoprecipitates. **(F)** Immunoblot validation of mTERF9 interactants identified by co-IP/MS analysis in absence of RNase. Replicate blots were probed with anti-Myc, anti-RH3, anti-CPN60α1/β1, anti-RPS1 and anti-RPL33. A replicate of a CBB-stained membrane is shown as input loading control.

To validate the mTERF9 interactome and its link with ribosome biogenesis, we performed immunoblot analyses of untreated mTERF9 co-immunoprecipitates and confirmed mTERF9 interaction with RH3, a DEAD box RNA helicase involved in rRNA processing (47), RPL33, RPS1 and the two chloroplast chaperonins CPN60α1 and β1 (Figure 7F).

In summary, the mTERF9 protein interactome is in agreement with the function of mTERF9 in chloroplast ribosome assembly. Moreover, the results demonstrate that mTERF9 protein supports protein-protein interaction during ribosome assembly besides its association with the *16S* rRNA. Finally, the striking interaction of CPN60 chaperonins with mTERF9 *in vivo* points towards the potential implication of the CPN60 complex in chloroplast translation.

### mTERF9 supports direct protein-protein and protein-RNA interactions

mTERF-repeat proteins are considered to be putative nucleic acid binders and we indeed showed that mTERF9 interacts with the *16S* rRNA *in vivo*. Moreover, our co-immunoprecipitation assays showed that mTERF9 interacts *in vivo* with many proteins that are involved in chloroplast ribosome biogenesis including ribosomal proteins, rRNA processing factors, and unexpected chaperonins from the CPN60 family. Some of these interactions appeared to be RNase insensitive. Together, the protein- and RNA-mTERF9 interactomes indicate that mTERF9 could support both protein-protein and RNA-protein interactions. To test the first possibility, we used mTERF9 as a bait in a modified yeast two-hybrid assay based on split-ubiquitin, called “DUAL hunter” (56). As mTERF9 and many ribosomal proteins partially associate to chloroplast membranes (57,58), this system offered the flexibility to select both membrane and cytosolic protein interactions. We tested the physical interaction of mTERF9 with 9 protein candidates that co-immunoprecipitated with mTERF9 in the -RNase or +RNase condition only or in both conditions (Figure 8 and Supplementary Figure 4A). The expression of the mTERF9 bait and the 9 prey proteins in the yeast co-transformants were verified by immunoblotting with anti-LexA and anti-HA antibodies, respectively (Supplementary Figure 4B). Out of the 9 candidates tested, mTERF9 interacted with 5 proteins (Figure 8B). These included ERA1, the Arabidopsis ortholog of the bacterial YqeH/ERA assembly factor for the 30S ribosomal subunit (59,60), PSRP2 and RPL1, two proteins of the 30S and 50S ribosomal subunits (61), respectively, and finally, CPN60β1 and β3, two subunits of the CPN60 chaperonin complex (62). These results demonstrated that mTERF9 can directly interact with proteins. The facts that mTERF9 interacts physically with ERA1, a protein that was specifically co-immunoprecipitated by the RNase treatment and with PSRP2 and RPL1 whose *in vivo* association with mTERF9 was rather sensitive to RNase, reinforced the notion that mTERF9 is likely to sequentially engage in various protein interactions during chloroplast ribosomal assembly and that the rRNAs likely stabilize some of these interactions. The physical interactions between mTERF9 and CPN60 chaperonins revealed that mTERF9 might be a substrate of the CPN60 complex. Alternatively, mTERF9 might recruit the CPN60 complex to ribosomal complexes to assist folding of ribosomal proteins during subunits assembly or neosynthesized proteins during translation. Finally, we demonstrated that mTERF9 had the capacity to self-interact in yeast and that the protein oligomerization was dependent on the mTERF repeats since their truncation abolished the interaction (Figure 8B and Supplementary Figure 4).

**Figure 8.**
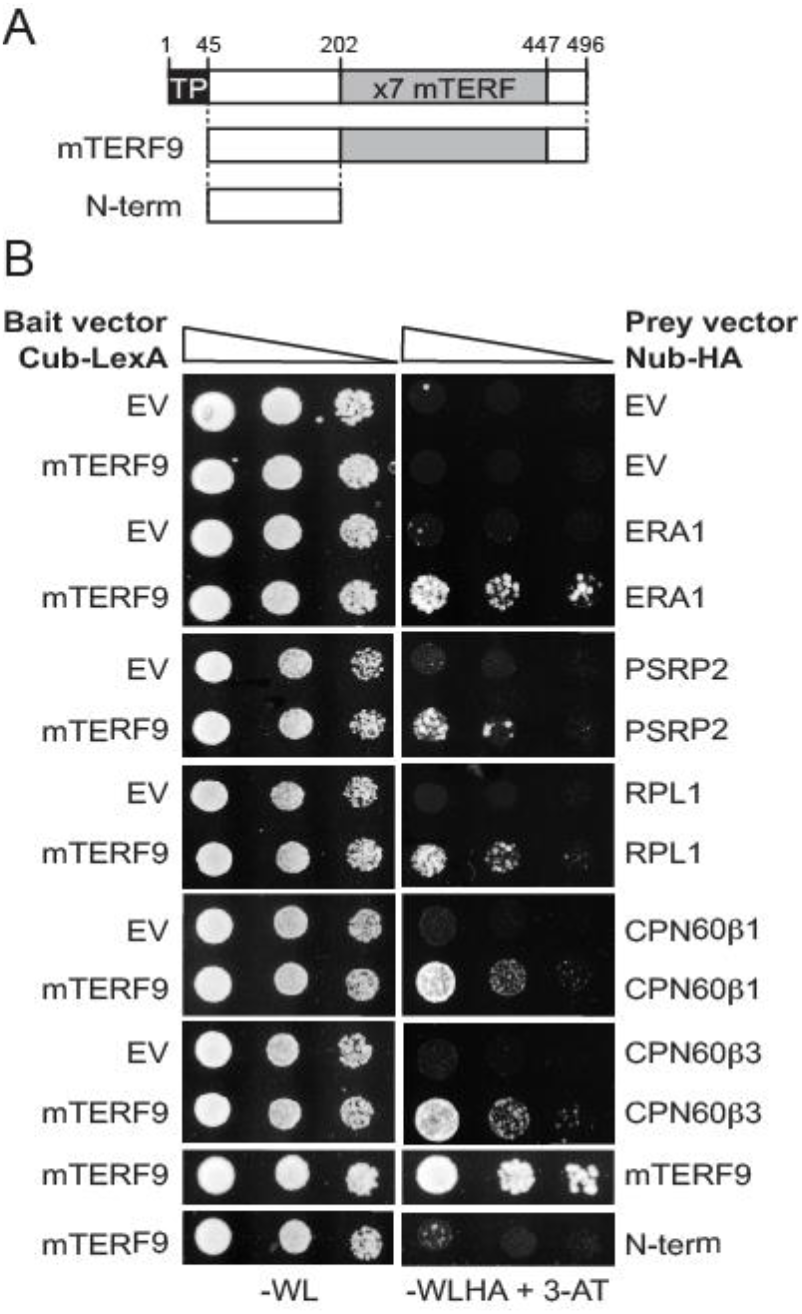
mTERF9 directly interacts with some of its *in vivo* protein interactants. **(A)** Schematic representation of mTERF9 used as bait or prey in the yeast two hybrid assay. **(B)** The yeast two hybrid assay was applied to assess direct interactions of mTERF9 with proteins identified by co-IP/MS analysis and mTERF9 self-association. mTERF9 interacts in yeast with ERA1, a putative 30S ribosomal subunit assembly factor, PSRP2 and RPL1, two plastid ribosomal proteins of the small and large ribosome subunits, CPN60β1 and β3, two subunits of the CPN60 chaperonin complex and finally itself. The bait vector expressing mTERF9 in fusion with the C-terminal half of the ubiquitin and the transcription factor LexA (Cub-LexA) was co-transformed with prey vectors expressing the protein candidates fused to the N-terminal half of the ubiquitin and HA tag (Nub-HA) in a yeast reporter strain. Yeast co-transformants were spotted in 10-fold serial dilutions on plates without Trp, Leu (-WL). Positive interactions allow growth on plates without Trp, Leu, His, Ade in presence of 3-aminotriazol (-WLHA + 3-AT). Negative controls were performed using bait or prey empty vectors (EV).

In a second time, we tested the capacity of mTERF9 to directly bind the *16S* rRNA using *in vitro* protein-RNA interaction assays. To this end, we expressed and purified *E. coli* recombinant mTERF9 (rmTERF9) fused to a maltose-binding protein (MBP) tag. The purity of the soluble rmTERF9 was visualized on SDS-PAGE and Coomassie Brilliant Blue staining (Figure 9A). A band migrated at the protein expected size (~96 kDa) but despite several attempts to optimize the purity of rmTERF9, a protein contaminant of ~60 kDa constantly copurified with rmTERF9. LC-MS/MS analysis confirmed the identity of rmTERF9 in the ~96 kDa band and identified *E. coli* GroEL, the ortholog of the Arabidopsis chloroplast CPN60 chaperonins, as the 60 kDa protein contaminant (Figure 9A). This result provides additional evidence for the direct interaction between mTERF9 and the CPN60 chaperonin complex. The purity of rmTERF9 was not adequate to detect specific RNA interactions by electromobility gel shift assay and, we therefore opted for the northwestern blot technique. This assay detects direct interaction between RNA and proteins that are immobilized on a membrane after their resolution by gel electrophoresis according to their charge and size (63,64), allowing the specific detection of rmTERF9 activity. The binding of mTERF9 to its *in vivo* target, the *16S* rRNA was compared to that for a chloroplast RNA of similar size from the *psbC* gene (Figure 9B and 9C). An interaction was detected between rmTERF9 and the *16S* rRNA but no binding activity was observed for GroEL nor the purified MBP (Figure 9C), indicating that it is mTERF9 moiety that harbors the RNA binding activity. In addition, no RNA binding activity could be detected for rMDA1, a DNA-binding mTERF protein, that promotes transcription in Arabidopsis (21). In contrast to the *16S* rRNA, at similar protein amounts, only residual binding was observed for the *psbC* RNA confirming that mTERF9 preferentially interacts with the *16S* rRNA. The *16S* rRNA is predicted to fold into four distinct domains (65,66) (Figure 9D) and we further explored mTERF9 binding specificity for each of the subdomains (Figure 9C). A binding activity was detected for the *16S* rRNA domains I, II and III but not for domain IV. At similar protein amounts, the binding of rmTERF9 was higher for domains I and II than for domain III suggesting its binding preference for these two RNA segments. In addition, rmTERF9 showed minimal binding to an unrelated *atpH* transcript of similar size than domains I-III confirming the protein specific binding towards the *16S* rRNA. Altogether, our results demonstrated that rmTERF9 is an RNA binding protein that preferentially binds to the *16S* rRNA, which is consistent with the *in vivo* data and the protein association with the 30S ribosomal subunit.

**Figure 9.**
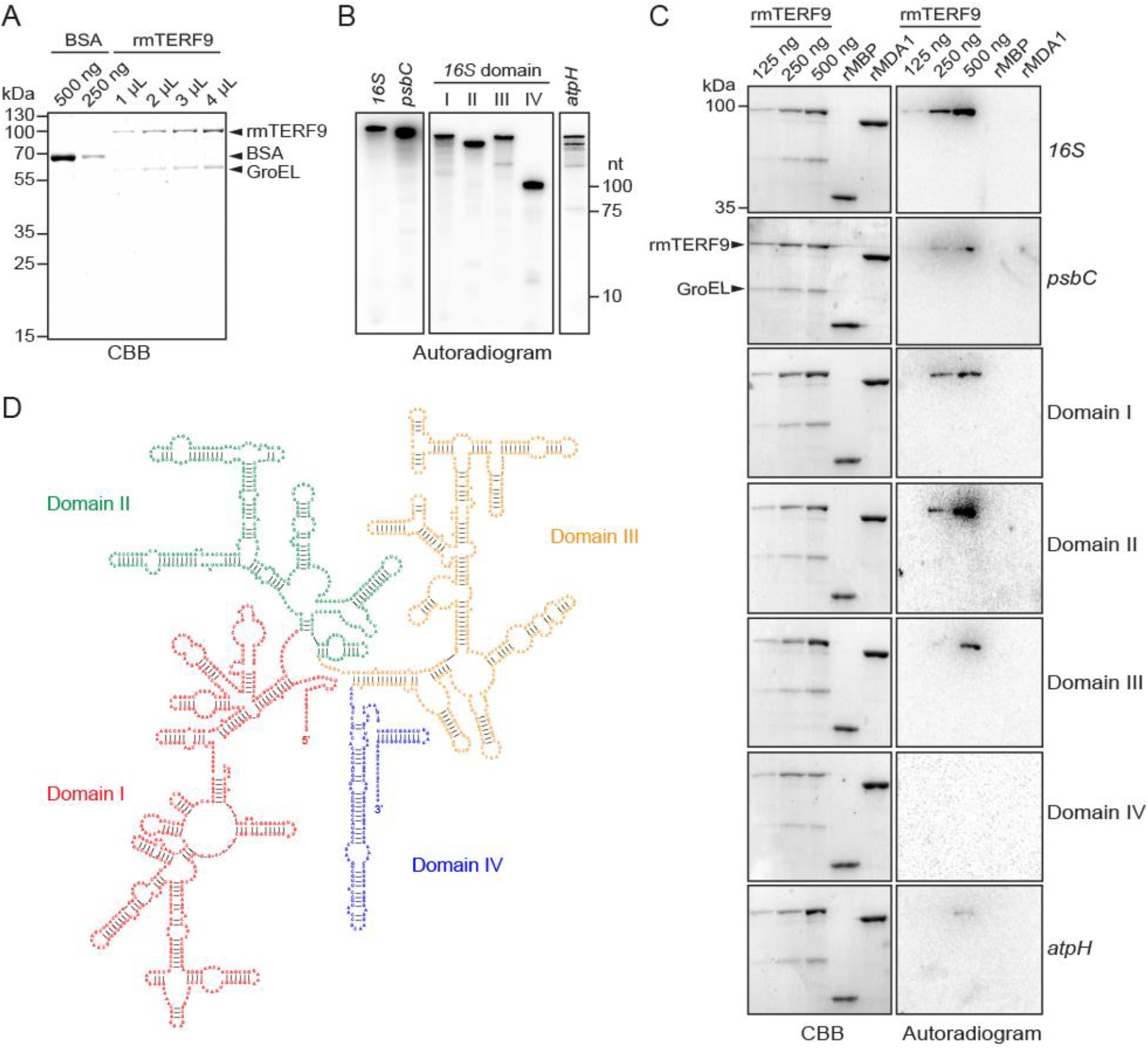
mTERF9 is an RNA binding protein that preferentially interacts with the *16S* rRNA *in vitro*. **(A)** Purification of rmTERF9. Increasing volumes of the purified rmTERF9 fraction were analyzed along with a BSA standard by SDS-PAGE and staining with CBB. **(B)** One μL of radiolabeled *in vitro* transcribed RNAs used in the northwestern assays were electrophoresed on a 7.5% denaturing polyacrylamide gel and autoradiographed. The RNA sizes are: *16S* rRNA, 1490 nt; *psbC*, 1422 nt; domain I, 508 nt; II, 356 nt; III, 478 nt; IV, 148 nt and *atpH*, 400 nt. **(C)** RNA binding activity of rmTERF9. The direct interaction between rmTERF9 and RNAs was tested by northwestern blotting. Increasing amount of rmTERF9 were resolved by SDS-PAGE and transferred to a PVDF membrane before hybridization with 0.5 fmole of radiolabeled RNA for *16S* and *psbC* or 1 fmole for domains I-V and *atpH*. rMBP and rMDA1 (500 ng each) were included as negative controls to show mTERF9 specific RNA binding activity. CBB stained membranes and autoradiograms are shown to the left and right, respectively. **(D)** Secondary structure of the Arabidopsis *16S* rRNA. The domains of the *16S* rRNA are labelled and delineated in color. The structure prediction is based on the secondary structure of bacterial *16S* rRNA (66) and was obtained from the RNACentral database (89).

## DISCUSSION

### mTERF9 assists chloroplast ribosome assembly via ribonucleoprotein interactions

We demonstrated in this study that mTERF9/TWIRT1, a member of the mTERF family of transcriptional factors in Arabidopsis has an unexpected function in chloroplast ribosome biogenesis and translation. Our study extends the current functional repertoire of mTERF proteins in plants in a process unrelated to DNA metabolism. We found that the *mterf9* knock-out line is defective in chloroplast translation as a result of the reduced accumulation of the *16S* and *23S* rRNAs, two scaffolding components of the 30S small and 50S large subunits of the chloroplast ribosome, respectively. The decrease of these rRNAs is intricately linked to the reduced assembly of functional chloroplast 70S ribosomes in *mterf9*. In fact, similar to bacteria, ribosome assembly in chloroplasts is tightly connected to the post-transcriptional maturation of rRNAs (67). For example, the orchestrated assembly of the 50S ribosomal proteins on the *23S* rRNA precursor in plants is believed to expose the RNA to endonucleases at particular cleavage sites and to generate the two hidden breaks in the *23S* rRNA. Consistent with that, the stability of several isoforms of the *23S* rRNA resulting from the hidden breaks processing were impaired in *mterf9*. Several auxiliary factors involved in chloroplast ribosomal assembly have been recently characterized and the majority of these are bacterial homologs or harbor RNA binding domains that are conserved in bacteria (47–51,55,68–70). Without any surprise, these protein homologs perform conserved functions in rRNA processing and therefore, ribosome assembly in chloroplasts. On the contrary, mTERF9 belongs to a eukaryote-specific transcription factors family and its function in chloroplast ribosome assembly was unexpected.

To firmly establish the *in vivo* function of mTERF9, we performed a comprehensive analysis of the *in vivo* RNA and protein interaction networks of mTERF9. Our co-immunoprecipitation results demonstrated that mTERF9 binds to the *16S* rRNA in chloroplasts as well as ribosomal proteins, CPN60 chaperonins and known auxiliary ribosomal factors involved in rRNA processing such as the MraW-like *16S* rRNA methyltransferase (CMAL) (69), YbeY endoribonuclease (68), RNase J (50), RNase E-like protein (RHON1) (51) or DEAD/DEAH-box RNA helicases (RH3 and ISE2) (47–49). The fractionation of chloroplast high-molecular-weight protein complexes combined with the comparative mTERF9 protein interactome in presence or absence of RNase together with the *16S* rRNA mTERF9 co-immunoprecipitation indicate that mTERF9 preferentially associates with the 30S small ribosome subunit *in vivo*. These results were highly consistent with the effects caused by the loss of mTERF9 in Arabidopsis, confirming the direct role of mTERF9 in chloroplast ribosomal assembly and translation.

Furthermore, we showed that some of the *in vivo* mTERF9 protein interactions could be reconstituted in a yeast two hybrid assay, demonstrating mTERF9 capacity to directly interact with ribosomal proteins. Besides supporting direct protein interactions, our RNA-protein interaction *in vitro* assays showed that mTERF9 is an RNA binding protein that preferentially binds the *16S* rRNA. The capacity to interact with proteins and/or nucleic acids are two potent biochemical properties that are intrinsically linked to α-helices structures (reviewed in 71,72), and the ability of mTERF9 to stabilize ribonucleoprotein complexes via physical interactions certainly accounts for its function in ribosomal assembly in chloroplasts. In addition, the capacity of mTERF9 to oligomerize via intermolecular interactions between the mTERF repeats likely confers the protein new opportunities for ligand association by extending the binding surfaces at the dimer (Figure 8B). Finally, mTERF9 association with the polysomes indicates that it plays a function in chloroplast translation after its initiation but how it participates in this process remains elusive at this stage (one possibility is discussed below).

### ERA1 and CPN60 chaperonins associate to chloroplast ribosomes *in vivo*

Our work revealed the presence of proteins in mTERF9 co-immunoprecipitates whose interaction with the chloroplast ribosomes had not been reported so far. The Arabidopsis protein ERA1 has been named after its bacterial homolog, the GTP-binding ERA protein. The protein localizes to the chloroplast nucleoids (73) and was found in stromal megadalton complexes containing ribosomal proteins (74) but its function has never been investigated in plants. In bacteria, the ERA1 homolog associates with the ribosome and binds the *16S* rRNA to promote the assembly of the small 30S subunit (59,60,75). Our results confirmed the physical interaction of Arabidopsis ERA1 with mTERF9, a protein involved in the assembly of the 30S ribosomal subunit and therefore, its likely conserved function in ribosome assembly in chloroplasts.

Another surprise in the composition of the mTERF9 protein interactome is the high-enrichment of the six subunits of the multi-subunit CPN60 chaperonin complex, which is related to the bacterial GroEL protein folding machine and has been proposed to share a conserved function in protein quality control in chloroplasts by preventing aberrant protein folding and aggregation during their import into chloroplast or during their synthesis in chloroplasts (62). However, the evidence for such function in plants is scarce and very few ligands of the CPN60 chaperonins have been reported so far. These include the large subunit of Rubisco, RbcL (76), the Ferredoxin NADP+ reductase, FNR (77), the NdhH subunit of the NADH dehydrogenase complex (78), the membrane-bound FTSH11 protease (79) and the Plastidic type I signal peptidase 1, Plsp1 (80). By confirming the *in vivo* association of the CPN60 complex with the chloroplast ribosomes and the direct interaction of CPN60β1 and β3 subunits with mTERF9, our study provides insights into the molecular function of these chaperonins in chloroplast translation. Based on the mTERF9 protein interactome and mTERF9’s *in vivo* function, we propose the CPN60 chaperonin complex to be involved in the folding of nascent chloroplast proteins during translation, which would be in agreement with the chaperonin paradigm (62). In this model, mTERF9 would serve as a platform for recruiting the CPN60 chaperonin complex to the chloroplast ribosomes during translation via direct protein-protein interactions. This assumption is tempting as it would explain the functional association of mTERF9 with the chloroplast polysomes besides its role in ribosome assembly. Alternatively, the physical interaction between mTERF9 and the CPN60 complex might reflect the direct involvement of CPN60 chaperonins in the folding of mTERF9 and/or ribosomal proteins during ribosome assembly. In this scenario, the CPN60 complex would alternatively play a direct role in chloroplast ribosome biogenesis and translation. This possibility has been foreshadowed in an early study that identified nuclear mutants of maize displaying defects in the assembly of chloroplast polysomes (81). The results showed that the product of a nuclear gene, *CPS2* facilitated the translation of various chloroplast mRNAs and the gene was later identified to be the maize orthologous *CPN60α1* gene (82).

### mTERF proteins as regulators of organellar translation

In metazoans, two out of the four mitochondrial mTERF proteins have been reported to regulate mitochondrial biogenesis and translation. The mTERF3 and mTERF4 proteins both interacts with mitochondrial rRNAs and are required for ribosomal assembly in mitochondria and therefore, translation (83,84). In addition, mTERF4 was shown to directly recruits the 5-methylcytosine methyltransferase, NSUN4 to the large ribosomal subunit to facilitate monosome assembly in mitochondria (84–86). Similar to our observations for *mterf9* in *Arabidopsis*, the loss of organellar translation in *mterf3* or *mterf4* mutants in mice led to an increase in the steady-state levels of mitochondrial transcripts and *de novo* transcription which were considered to be a secondary effect of the loss of mitochondrial translation (84,85). On contrary in plants, mTERF9 is so far the only mTERF protein reported to play a direct role in ribosomal assembly and chloroplast translation (87). Two other members, mTERF4 and mTERF6 in maize and Arabidopsis respectively, have been reported to influence chloroplast translation but their effect on translation was rather indirect (22,44). In fact, mTERF4 promotes the splicing of several RNAs encoding ribosome components whereas mTERF6 contributes to the maturation of the *trnI.2* in chloroplasts. As a consequence, the loss of function of these proteins caused a reduced accumulation of chloroplast ribosomes and translation in plants. By contrast, our comprehensive analysis of mTERF9 function that combined reverse genetics, molecular and biochemical phenotyping as well as *in vitro* assays allowed us to make firm conclusion about mTERF9 implication in chloroplast ribosome biogenesis. Our study demonstrated that mTERF9 supports physical *in vivo* association with RNA and protein components of the ribosome to stimulate ribosomal assembly and chloroplast translation, confirming the conserved function of mTERF-repeat proteins in the regulation of organellar translation in the plant kingdom.

## SUPPLEMENTARY DATA

Supplementary Data are available at NAR online: Supplementary Figure 1, Supplementary Figure 2, Supplementary Figure 3, Supplementary Figure 4, Supplementary Table 1, Supplementary Data Set 1, and Supplementary Data Set 2.

**Supplementary Figure 1.**
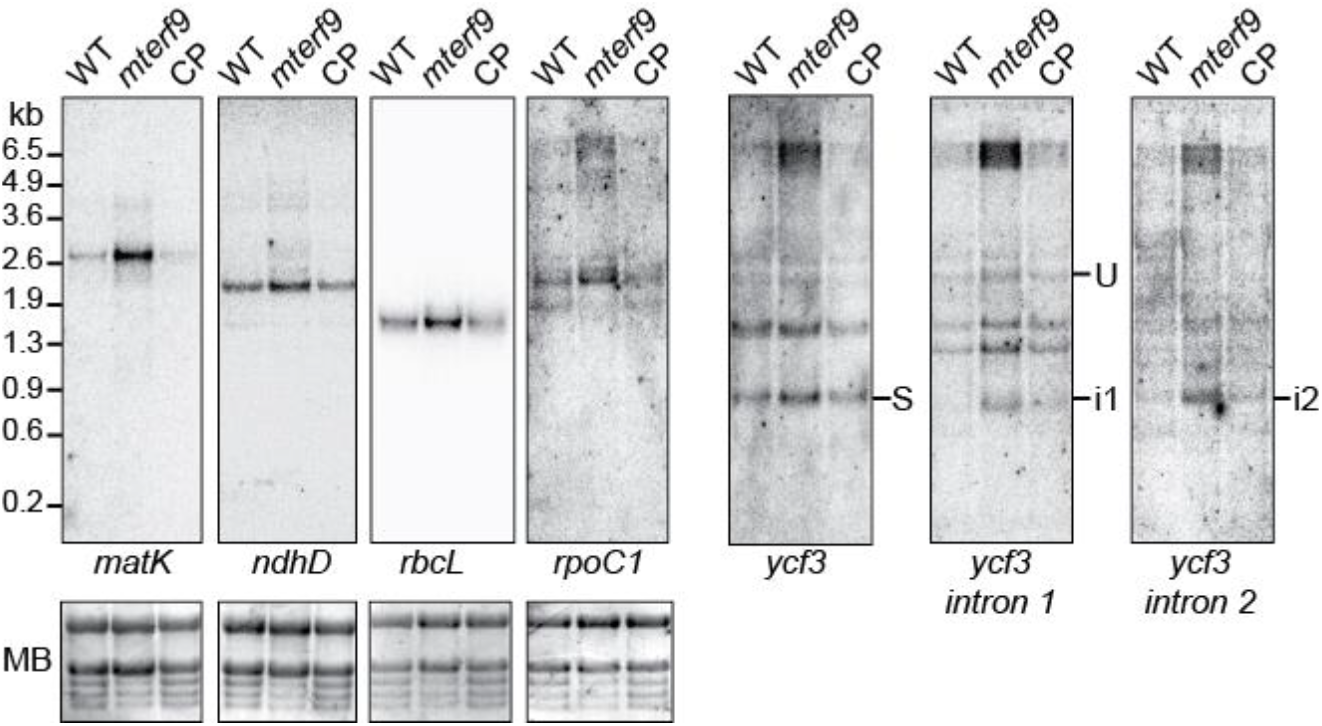
Northern blot analysis of selected chloroplast gene in WT, *mterf9* and CP plants. An excerpt of the methylene blue stained blots is shown to illustrate equal loading. *matK*, *ndhD*, *rbcL* blots were striped before rehybridization with *ycf3*, *ycf3-1*, and *ycf3-2* probes, respectively.

**Supplementary Figure 2.**
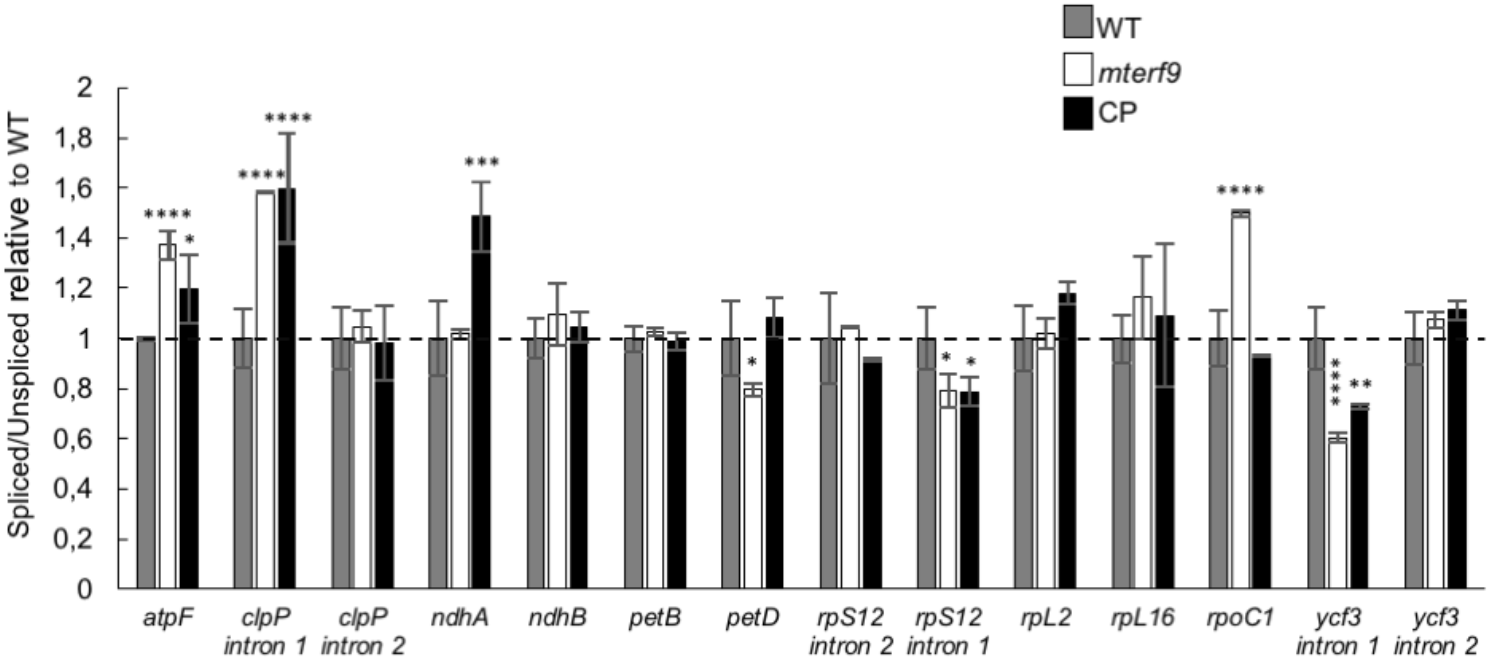
Chloroplast RNA intron splicing efficiency in WT, *mterf9* and CP plants. ANOVA, Dunnet’s multiple test correction, *, *P* < 0.05.

**Supplementary Figure 3.**
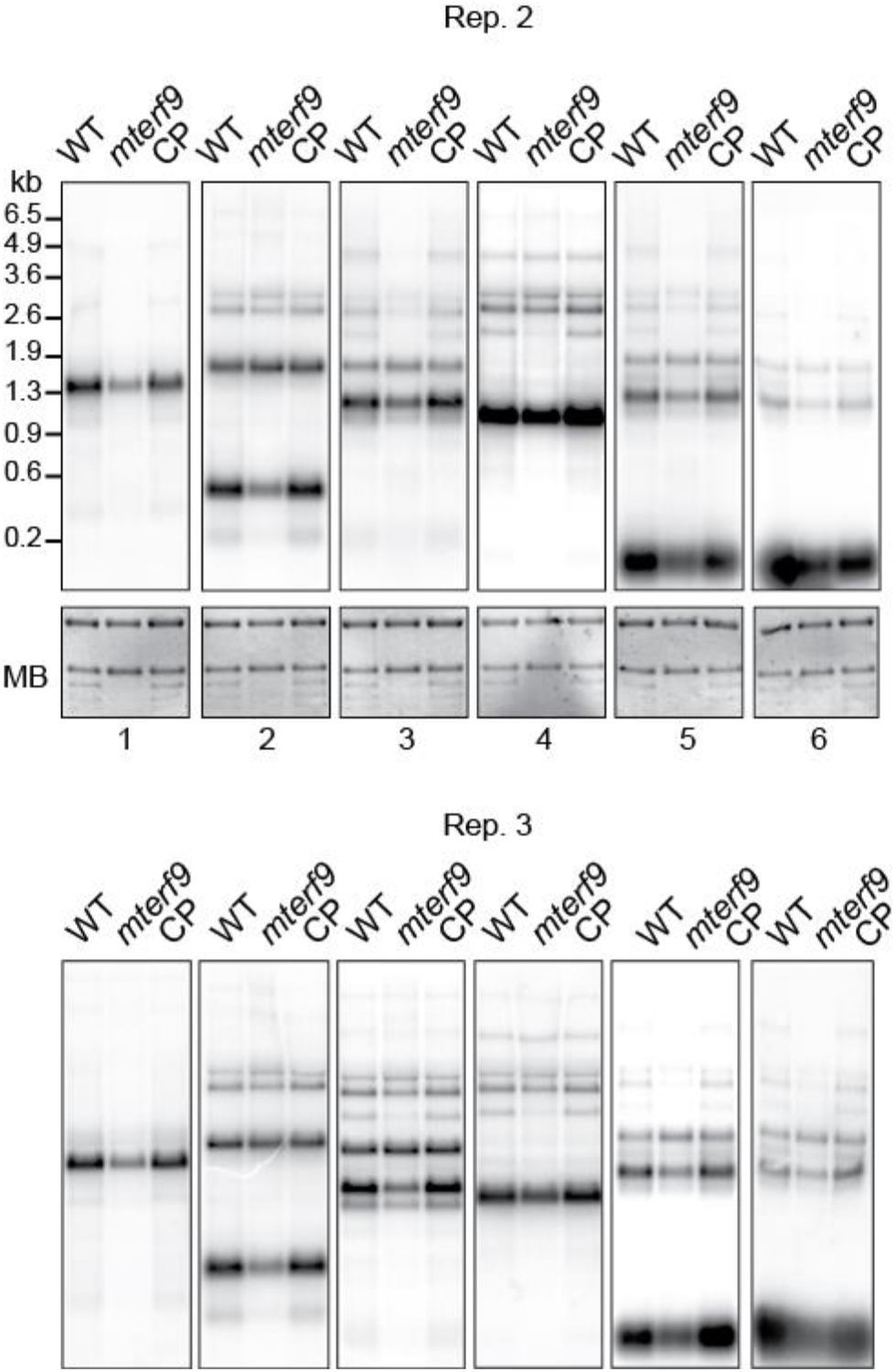
Chloroplast rRNA northern blot replicates in WT, *mterf9* and CP plants. In support of Figure 4.

**Supplementary Figure 4.**
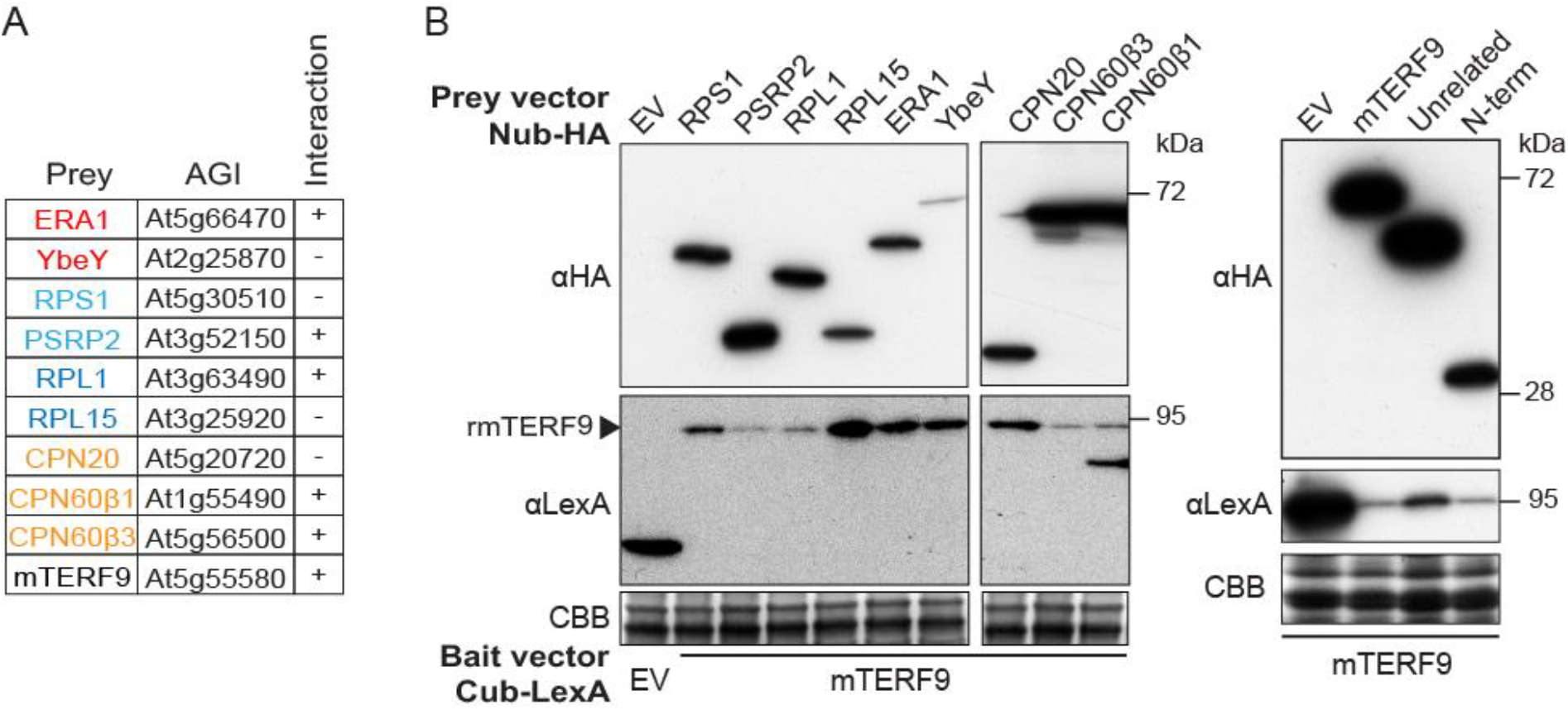
**(A)** Prey and bait interactions tested in the yeast two hybrid assays. **(B)** Immunoblot analysis of prey and bait proteins expression in yeast. Total proteins were fractionated by SDS-PAGE. The CBB stained membrane serves as a loading control. The bait and prey proteins were detected using anti-LexA and -HA antibodies, respectively. EV: empty vector. The immunoblot to the right displays a sample in the middle lane that is unrelated to this study.

**Supplementary Table 1.**
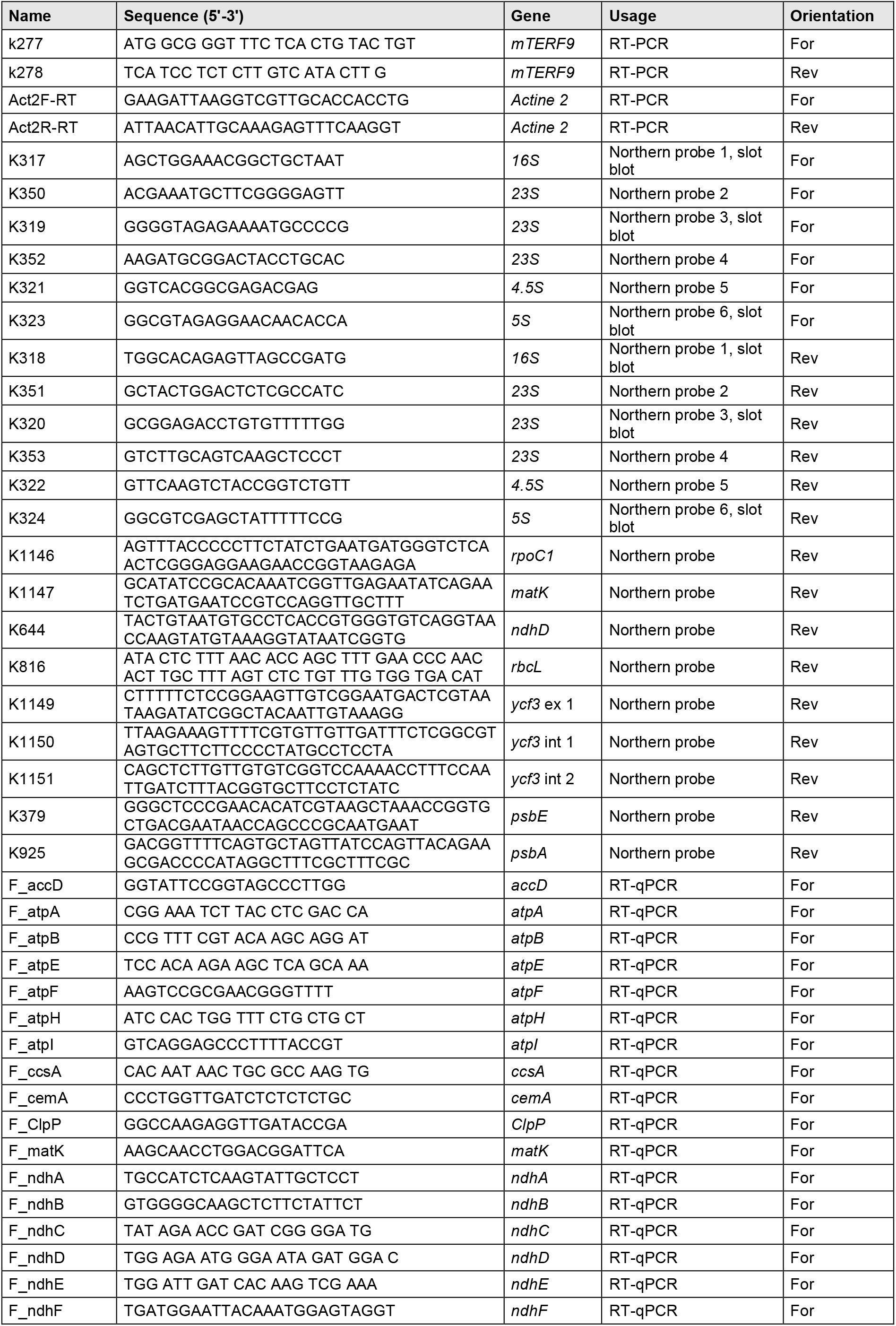
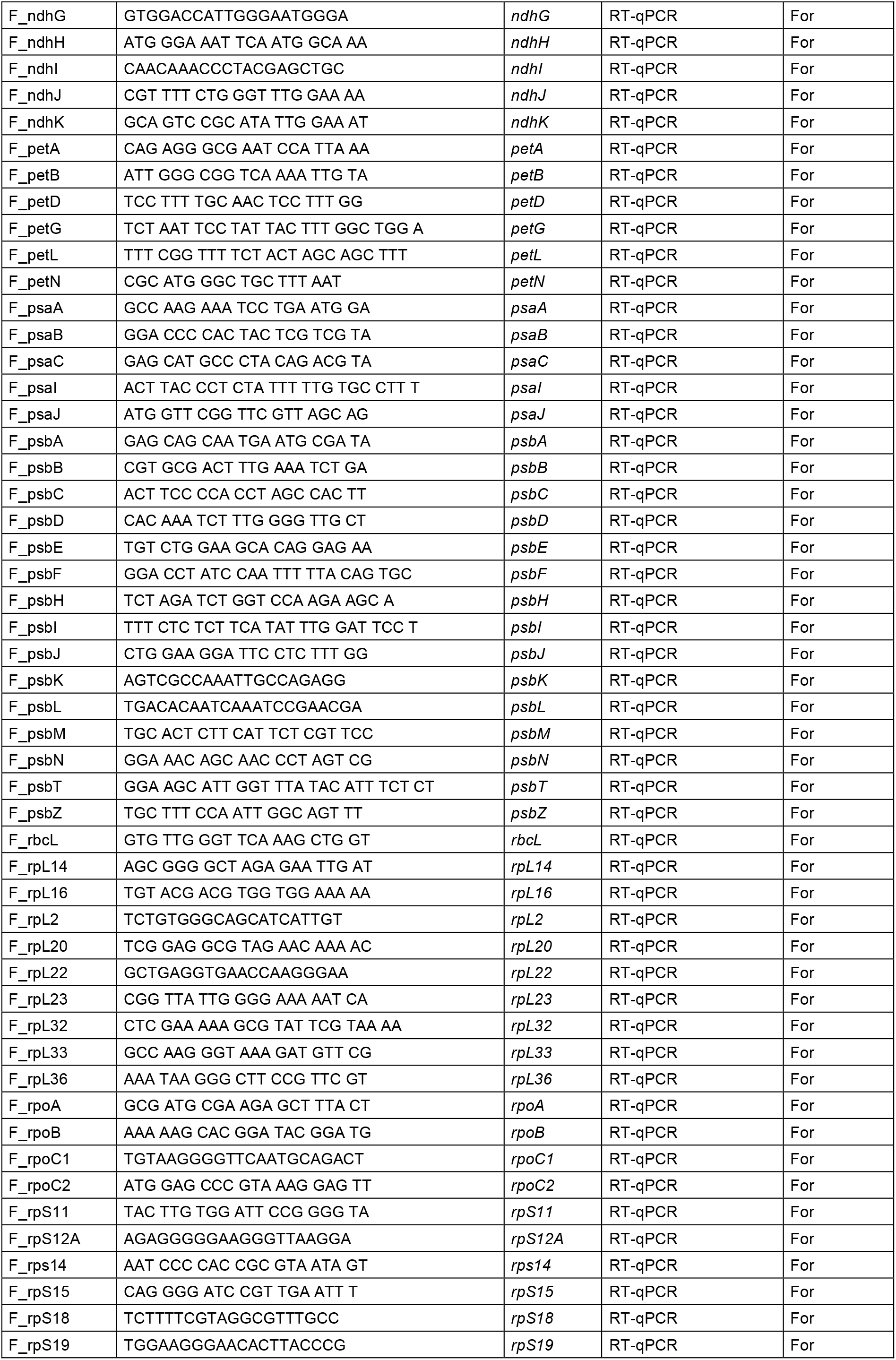
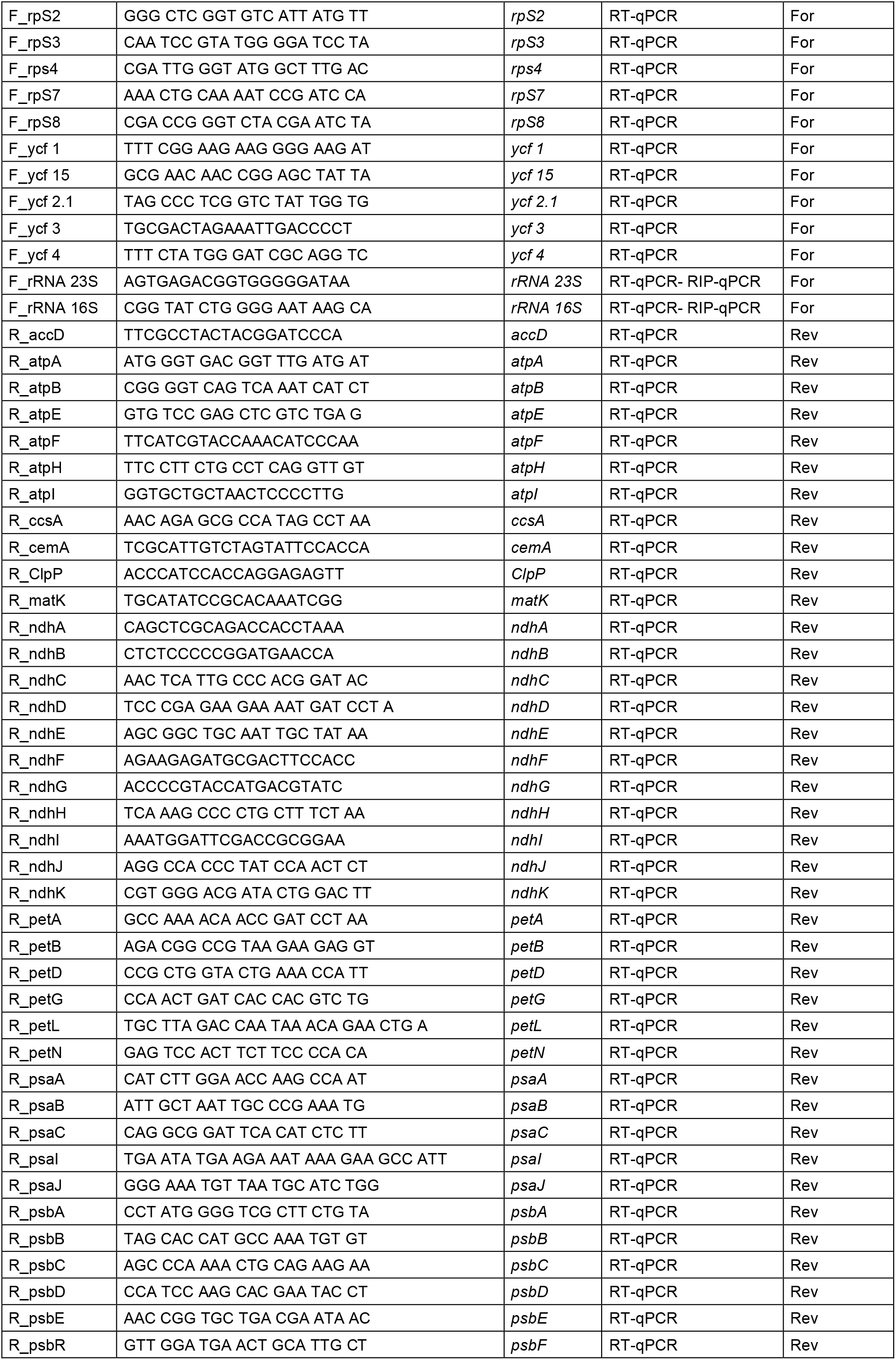
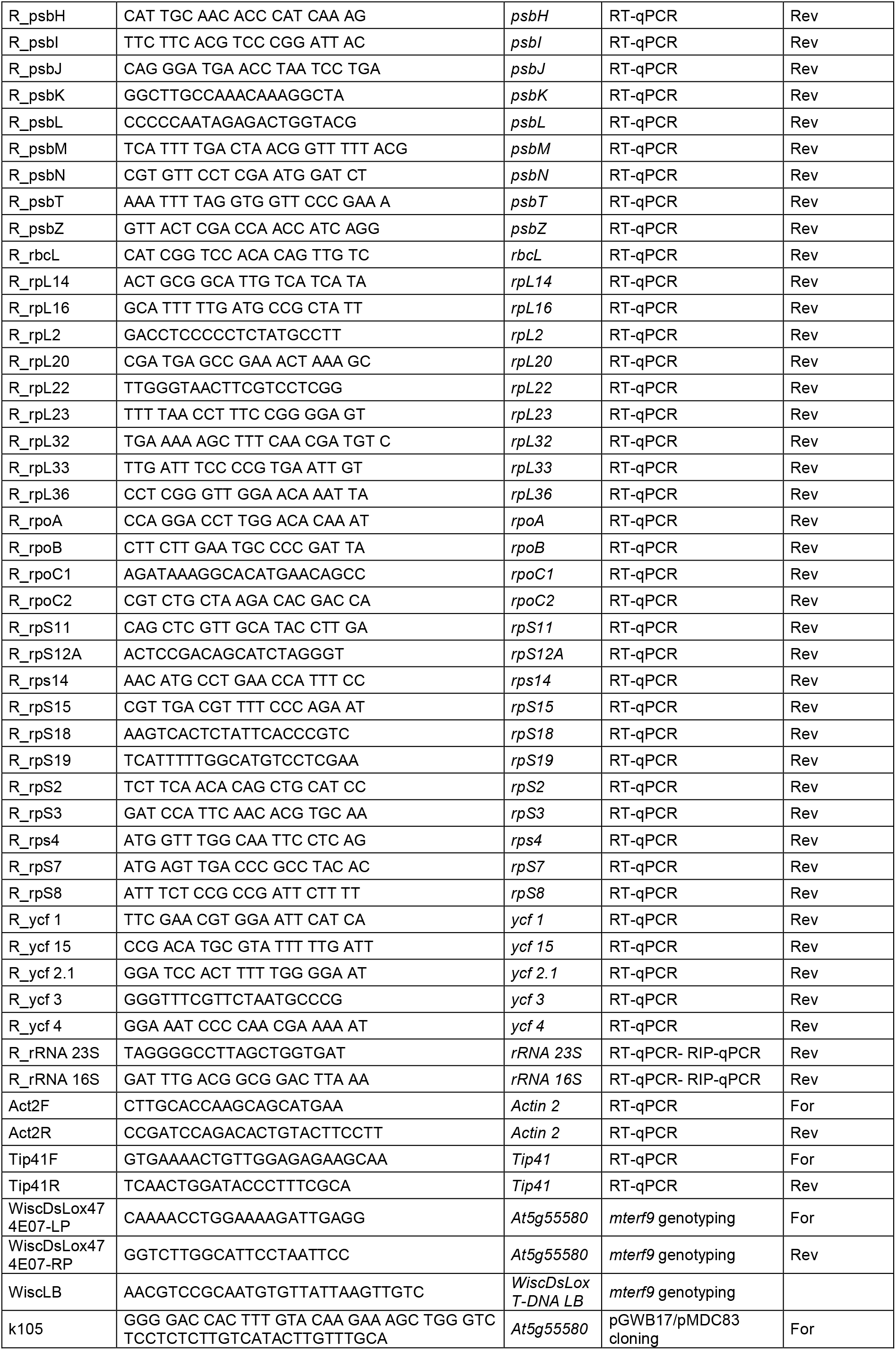
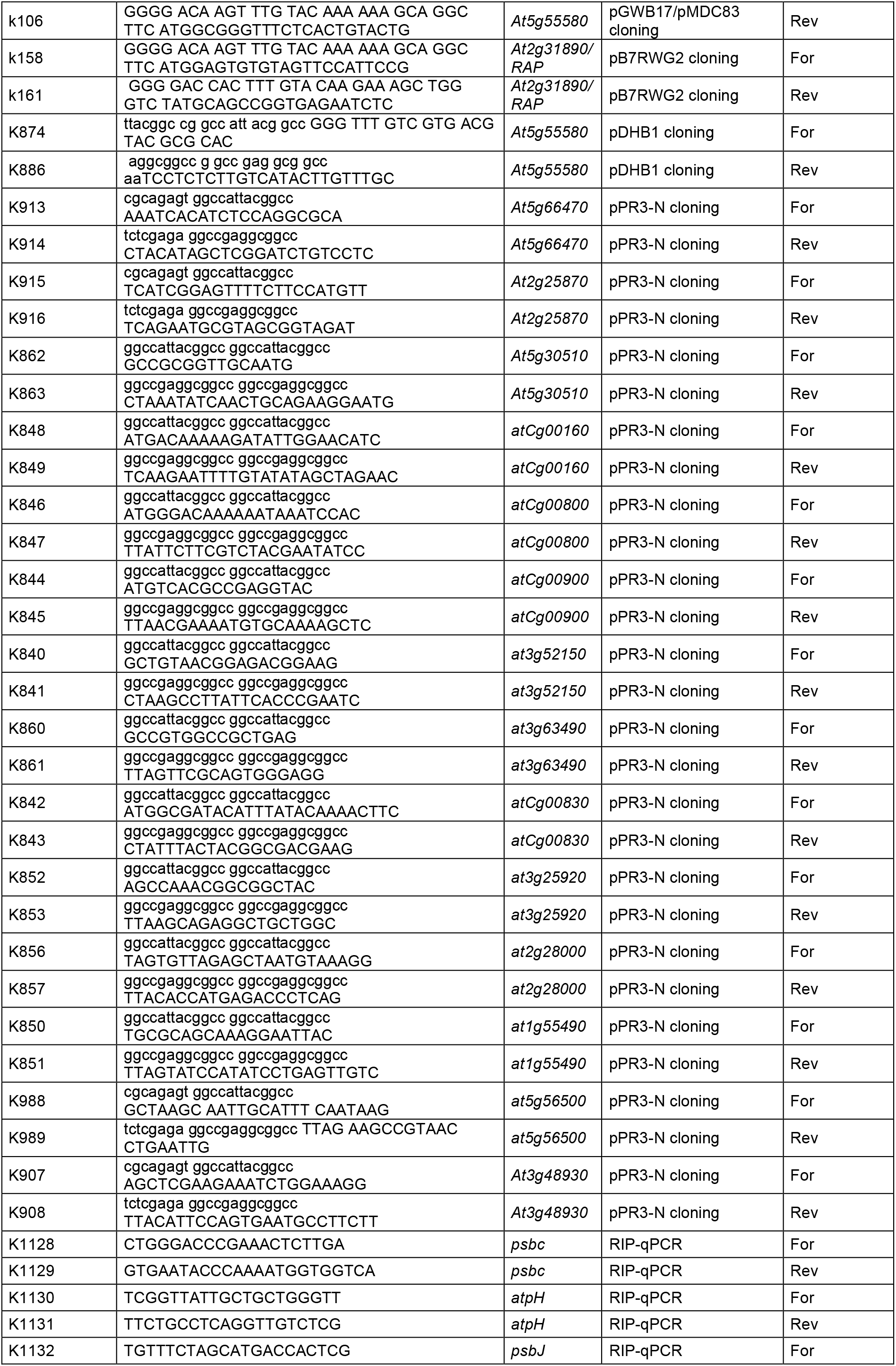
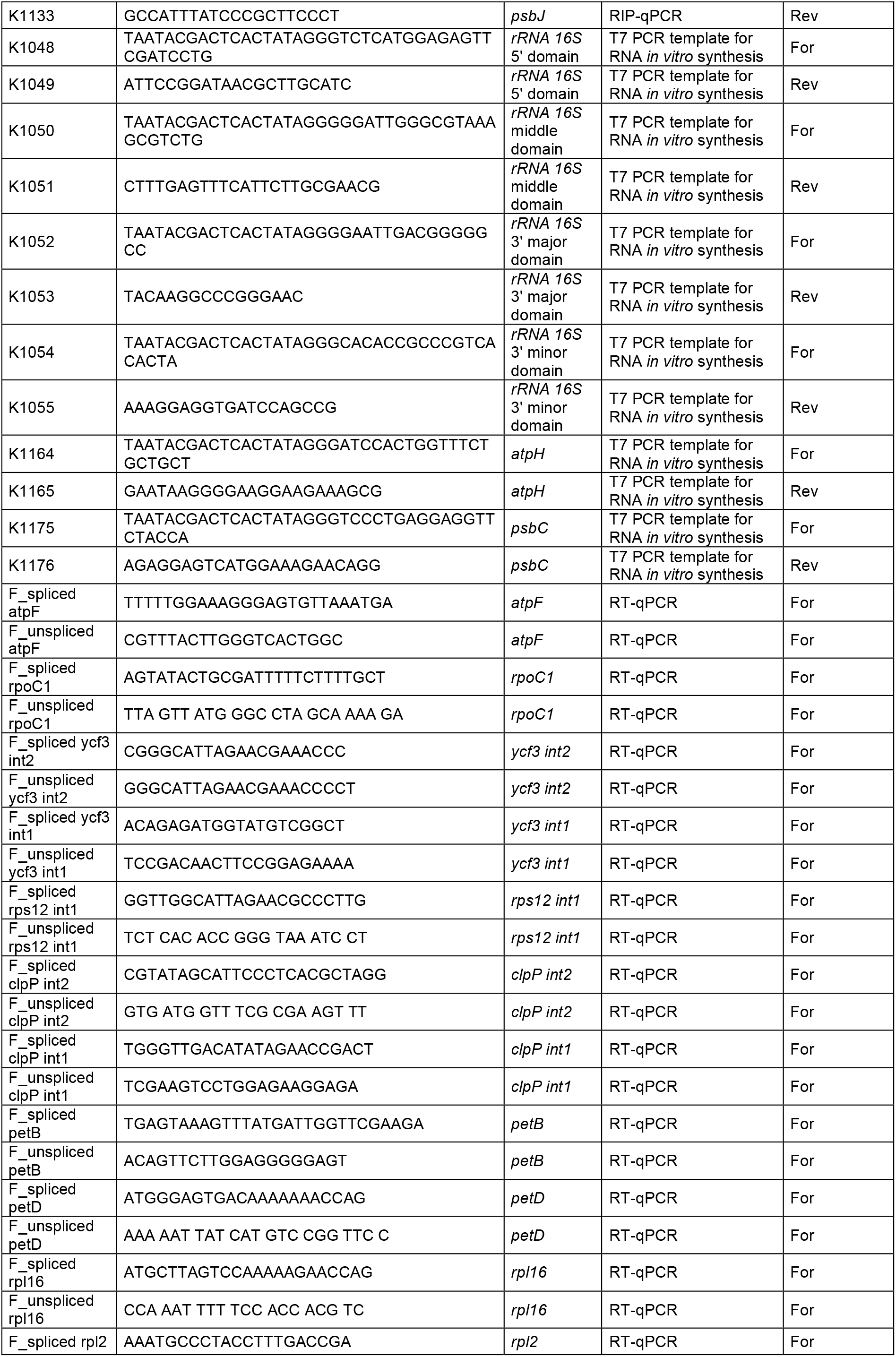
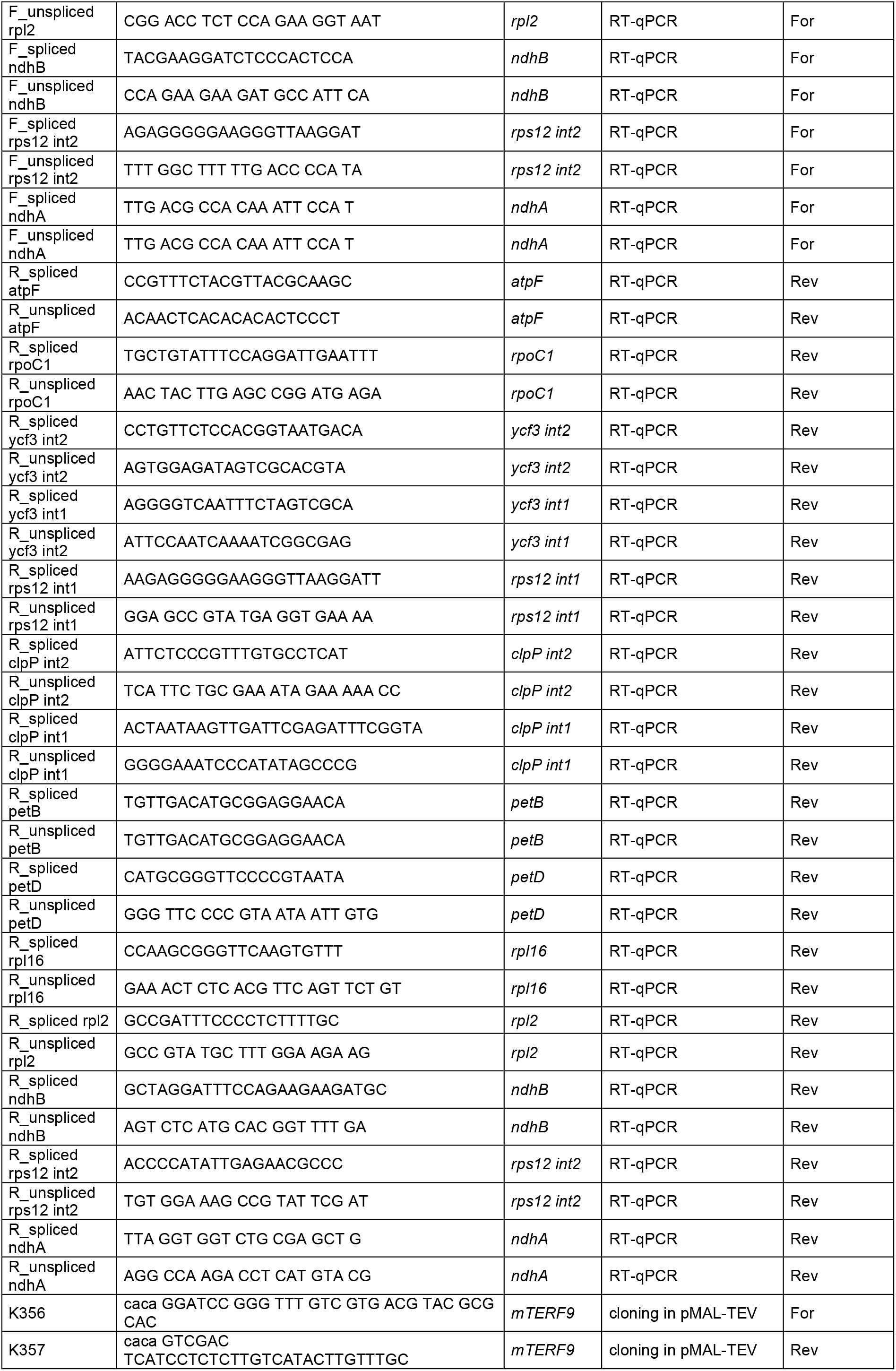
List of oligonucleotides used in this study.

**Supplementary Data Set 1.** List of proteins identified by LC-MS/MS in co-immunopurification assays using mTERF9 as bait.

**Supplementary Data Set 2**. RIP-Chip data showing enrichment of RNA sequences in mTERF9 immunoprecipitations.

